# Protein phosphatase 1 regulates core PCP signaling

**DOI:** 10.1101/2023.09.12.556998

**Authors:** Song Song, Bomsoo Cho, Alexis T. Weiner, Silas Boye Nissen, Irene Ojeda Naharros, Pablo Sanchez Bosch, Kaye Suyama, Yanhui Hu, Li He, Tanya Svinkina, Namrata D. Udeshi, Steven A. Carr, Norbert Perrimon, Jeffrey D. Axelrod

## Abstract

PCP signaling polarizes epithelial cells within the plane of an epithelium. Core PCP signaling components adopt asymmetric subcellular localizations within cells to both polarize and coordinate polarity between cells. Achieving subcellular asymmetry requires additional effectors, including some mediating post-translational modifications of core components. Identification of such proteins is challenging due to pleiotropy. We used mass spectrometry-based proximity labeling proteomics to identify such regulators in the *Drosophila* wing. We identified the catalytic subunit of Protein Phosphatase1, Pp1-87B, and show that it regulates core protein polarization. Pp1-87B interacts with the core protein Van Gogh and at least one Serine/Threonine kinase, Dco/CKIc, that is known to regulate PCP. Pp1-87B modulates Van Gogh subcellular localization and directs its dephosphorylation in vivo. PNUTS, a Pp1 regulatory subunit, also modulates PCP. While the direct substrate(s) of Pp1-87B in control of PCP is not known, our data support the model that cycling between phosphorylated and unphosphorylated forms of one or more core PCP components may regulate acquisition of asymmetry. Finally, our screen serves as a resource for identifying additional regulators of PCP signaling.

## Introduction

PCP signaling controls the polarization of cells, typically within the plane of an epithelium, orienting asymmetric cellular structures, cell divisions and cell migration. Much mechanistic understanding of PCP signaling has and will likely continue to derive from work using *Drosophila* as a model system. In flies, PCP signaling controls the orientation of hairs on the adult cuticle, chirality and orientation of ommatidia in the eye, orientation of cell divisions, and related processes in other tissues(Butler & Wallingford, 2017; Harrison *et al*, 2020). While much has been learned from studies in flies, medically important developmental defects and physiological processes in vertebrates are also under control of PCP signaling. Defects in the core PCP mechanism result in a range of developmental anomalies and diseases including open neural tube defects (reviewed in (Copp & Greene; Simons & Mlodzik, 2008)), conotruncal heart defects (Garriock *et al*, 2005; Gibbs *et al*, 2016; Henderson Phillips & Chaudhry, 2006; Phillips *et al*, 2005; Phillips *et al*, 2007), deafness (reviewed in(Lu & Sipe, 2016; Montcouquiol & Kelley, 2020)), situs inversus and heterotaxy (reviewed in (Santos & Reiter)). PCP is also believed to participate in both early and late stages of cancer progression (reviewed in(VanderVorst *et al*, 2019; Zhan Rindtorff & Boutros, 2017)) and during wound healing (Caddy *et al*; Lee & Adler, 2004). PCP polarizes skin and hair (Devenport & Fuchs, 2008; Guo Hawkins & Nathans, 2004; Wang Badea & Nathans, 2006; Wang Guo & Nathans, 2006) and the ependyma (Guirao *et al*, 2010; Hirota *et al*, 2010; Ohata *et al*, 2014; Tissir *et al*, 2010). Much of our knowledge of PCP signaling derived from the *Drosophila* model translates to studies of PCP in vertebrate model systems, thus informing the pathogenesis of these conditions.

The dedicated components of a core signaling module that acts to generate and amplify molecular asymmetry, and to coordinate the direction of polarization between neighboring cells, have been recognized for some time. Six known proteins in the core signaling module, are the serpentine protein Frizzled (Fz) (Vinson & Adler, 1987; Vinson Conover & Adler, 1989), the seven-pass atypical cadherin Flamingo (Fmi; a.k.a. Starry night) (Chae *et al*, 1999; Usui *et al*, 1999), the 4-pass protein Van Gogh (Vang; a.k.a. Strabismus) (Taylor *et al*, 1998; Wolff & Rubin, 1998), and the cytosolic/peripheral membrane proteins Dishevelled (Dsh) (Klingensmith Nusse & Perrimon, 1994; Thiesen *et al*, 1994), Diego (Dgo) (Feiguin *et al*, 2001), and the PET/Lim domain protein Prickle (Pk)(Gubb *et al*, 1999), which adopt asymmetric subcellular localizations that predict the morphological polarity pattern such as hairs in the fly wing (reviewed in (Butler & Wallingford, 2017; Zallen, 2007)). These proteins communicate at cell boundaries, recruiting one group to the distal side of cells, and the other to the proximal side, through the function of an incompletely understood feedback mechanism, thereby aligning the polarity of adjacent cells. The direction of core module polarization is proposed to be responsive to several kinds of signals tied to the tissue axes, including expression gradients, diffusible ligands and mechanical forces (reviewed in (Butler & Wallingford, 2017; Eaton & Julicher, 2011)).

A variety of post-translational modifications to the core PCP components have been recognized as regulators of polarization, and include serine-threonine and tyrosine phosphorylation, farnesylation, and ubiquitinylation (reviewed in(Harrison *et al*., 2020)). Furthermore, processes including oligomerization, internalization and regulated transport are thought to play key roles in regulating polarization. Understanding these events is critical to understanding the PCP mechanism, yet a challenge has been to identify the proteins that carry them out, as many are broadly pleiotropic and are therefore not easily recognized in genetic screens. We report here the results of an APEX based proximity labeling screen. APEX, an active peroxidase, biotinylates proteins within a 5-50nm radius of the probe in living cells(Bosch Chen & Perrimon, 2021). This approach has several times been used successfully in living *Drosophila* tissues(Chen *et al*, 2015; Li *et al*, 2020; Liu *et al*, 2020; Mannix *et al*, 2019). By fusing APEX to the core PCP protein Vang, we labeled proteins likely to interact with the core PCP proteins in living *Drosophila* wing discs. In a secondary RNAi screen, many of these proteins can be shown upon knockdown to produce either or both of two typical PCP polarity phenotypes: misoriented wing hairs or a multiple wing hair phenotype.

Because several serine/threonine kinases have been suggested to regulate core PCP function, from our candidate list, we chose to further characterize Pp1-87B, which encodes the catalytic subunit of protein phosphatase 1 (Pp1). By appropriately timed knockdown, we show that Pp1- 87B activity is necessary for properly oriented wing hairs, and for controlling the amount and subcellular localization of two core PCP proteins, Vang and Pk. Pp1-87B knockdown genetically interacts with core PCP mutants, and with Dco/CKIε;, the *Drosophila* and vertebrate homologs of a S/T kinase thought to regulate PCP in both *Drosophila* and vertebrates. We find that Pp1-87B, either directly or indirectly, induces dephosphorylation of Vang in vivo. Furthermore, we identify PNUTS as a possible regulatory subunit of Pp1 that may provide specificity for PCP. Furthermore, our potential PCP interactome will serve as a resource for identification of other regulators of PCP signaling.

## Results

### Proximity labeling screen

To identify proteins participating in PCP that are not readily discovered in genetic screens, we fused APEX, along with myc and V5 tags, to the N-terminus of Vang and expressed this under control of D174-GAL4 or 71B-GAL4, that each express throughout the wing. APEX::Vang expression in a *vang* mutant background showed normal planar polarity, verifying functionality of the engineered protein (**Expanded View Figure EV1**). Streptavidin staining of third instar wing discs fixed after labeling detected biotinylation primarily at apicolateral junctions, substantially co-localizing with Pk, similar to localization of native Vang. Furthermore, both Pk and Fmi were biotinylated by APEX::Vang *in vivo* (**Expanded View Figure EV2**). Detection of Pk and Fmi, both known to be present in proximal PCP complexes, gave us confidence that APEX::Vang could effectively label the proximal PCP proteome. We note, however, that during polarization, complexes of opposite orientation are thought to interact, and indeed Vang has been shown to physically Interact with each of the other core proteins(Adler, 2012; Goodrich & Strutt, 2011; Seifert & Mlodzik, 2007), so that we expect to also label proteins associated with the distal complex.

Since planar polarization is known to be underway in third instar(Sagner *et al*, 2012), we probed the PCP proteome in third instar wing discs. Four pools, two carrying D174-GAL4, UAS-APEX::Vang (specific) and two carrying only D174-GAL4 to be used for subtraction of background biotinylation (background), were collected and incubated with biotin-phenol in S2 medium. Biotinylated proteins from lysates were enriched on streptavidin beads, released from beads by on-bead trypsin digestion and labeled with iTRAQ (isobaric tags for relative and absolute quantification) 4-plex reagents followed by liquid chromatography-mass spectrometry (LC-MS/MS). The relative level of a given protein identified by at least two unique peptides in each sample (2028 in total; Expanded View Dataset 1) was obtained, and for each sample, the proteins in each background sample were used to generate the iTRAQ ratios. In total, four sets of iTRAQ ratios were generated (APEX::Vang replicate 1/control replicate 1; APEX::Vang replicate 1/control replicate 2; APEX::Vang replicate 2/control replicate 1; APEX::Vang replicate 2/control replicate 2). A false positive rate (FPR) cutoff was calculated based on the distribution of positive and negative control genes. The genes known and assumed positive or thought to be functional regulators of, or otherwise associated with, PCP are positive controls while those known to localize in compartments such as the nucleus where PCP is inactive, are negative controls(Rhee *et al*, 2013). At 10% FPR threshold, 123 proteins scored in three or four datasets and are reported as the high confidence candidate PCP interactome, and an additional total of 520 proteins scored in two of four datasets and were considered a low confidence interactome (Expanded View Dataset 1). As expected, the high priority candidate interactome included several known and likely PCP related components. Many of the identified proteins have known relationships to other proteins, as determined using COMPLEAT (protein COMPLex Enrichment Analysis Tool; http://www.flyrnai.org/compleat/), and when their related proteins were also identified, this is also shown. We prioritized the high confidence interactome list for a secondary RNAi screen, and to this list, we added proteins that scored below this cutoff, but that we deemed to be good candidates either because they co-complex with the highest scoring proteins or based on the nature of the gene product.

Using available RNAi lines from the TRiP and VDRC collections, we then performed a secondary RNAi screen for the candidate protein-coding genes to test for abnormal adult wing hair polarity phenotypes characteristic of known PCP gene mutants. We initially drove RNAi expression with *ptc*-Gal4 in the presence of Dicer-2, and for lines that produced lethality, we re-screened with clonal GAL4 expression and in some cases hh-GAL4, ci-GAL4 or ap-GAL4. We note that efficiency of somatic expression varies with the different VALIUM vectors in the TRiP collection, so that negative results, particularly for weaker expressors, may be false negatives (see materials and methods).

Expanded View Dataset 2 summarizes all of the proteins that scored positive in the RNAi screen for adult wing hair PCP phenotypes, specifically wing hair orientation defects most characteristic of core PCP mutants, and multiple wing hair phenotypes more characteristic of PCP effector mutants. Phenotypes were subjectively scored on a scale of 0-3 (no phenotype to strong phenotype), keeping in mind that a large contributor to strength is likely to be the specific properties of each individual RNAi line. This list comprises 91 proteins and includes 40 of the 123 high priority candidates from the APEX screen. A complete summary of the RNAi screen, including information about which RNAi lines were used, is shown in Expanded View Dataset 3. The data has been submitted to RSVP database, which is the resource for tracking and sharing the phenotype data of RNAi transgenic reagents (Perkins *et al*, 2015) (https://www.flyrnai.org/cgi-bin/RSVP_search.pl).

### Identification of Pp1-87B as a regulator of PCP signaling

Among the high priority screened proteins, we identified the protein phosphatase 1 catalytic subunit Pp1-87B. Three RNAi lines (BM32414, BM67911, VDRC35024) driven by *ptc*-gal4 produced considerable lethality, but escapers showed subtle wing hair orientation defects (**Appendix Figure S1**). The two Bloomington lines, BM32414 and BM67911, target independent sequences and by qRT-PCR suppress *Pp1-87B* mRNA by roughly half. Furthermore, expression of the Pp1-87B inhibitor I-2 expressed under *ptc*-gal4 control phenocopies that of the three RNAi lines, demonstrating that the observed polarity phenotypes are attributable to Pp1-87B-associated on target effects (**Appendix Figure S2**). Making RNAi expressing clones, we observed some swirls indicative of polarity defects. However, in pupal wings, a fluorescent clone marker showed dispersed signal, suggesting that some or all of the observed clonal phenotypes might result from the expected Pp1-87B-dependent cell lethality and resulting scars. Nine of the 122 proteins were co-complexed with Pp1-87B (Dataset 1; http://www.flyrnai.org/compleat/), suggesting possible relationships between Pp1-87B and other potential regulators of PCP. We note, however, that many substrates are expected, as Pp1-87B is thought to be responsible for 80% of the phosphatase activity in *Drosophila*(Dombradi *et al*, 1990).

To further evaluate a possible role for Pp1-87B in PCP control, we circumvented the lethality and other phenotypes associated with Pp1-87B knockdown in the *ptc* domain by using a temperature sensitive allele of GAL80 to restrict loss of function to the early pupal period during which wing hair polarity is established. Using RNAi line (32414), white prepupae were raised at the restrictive temperature (18°) before shifting to the permissive temperature (29°) at the equivalent of either 12 hr APF or 20 hr APF (**Figure 1**). Adult wings consistently showed abnormal hair orientation in the *ptc* domain both distal and proximal to the anterior crossvein when shifted to the restrictive temperature at 12 hr and much less frequently at 20 hr APF. Similarly, when expressed throughout the wing beginning using *actin-GAL4*, polarity perturbation is seen in other regions of the wing (**Expanded View Figure EV3**). Thus, appropriately timed loss-of-function produced consistent PCP hair orientation phenotypes.

**Figure 1.**
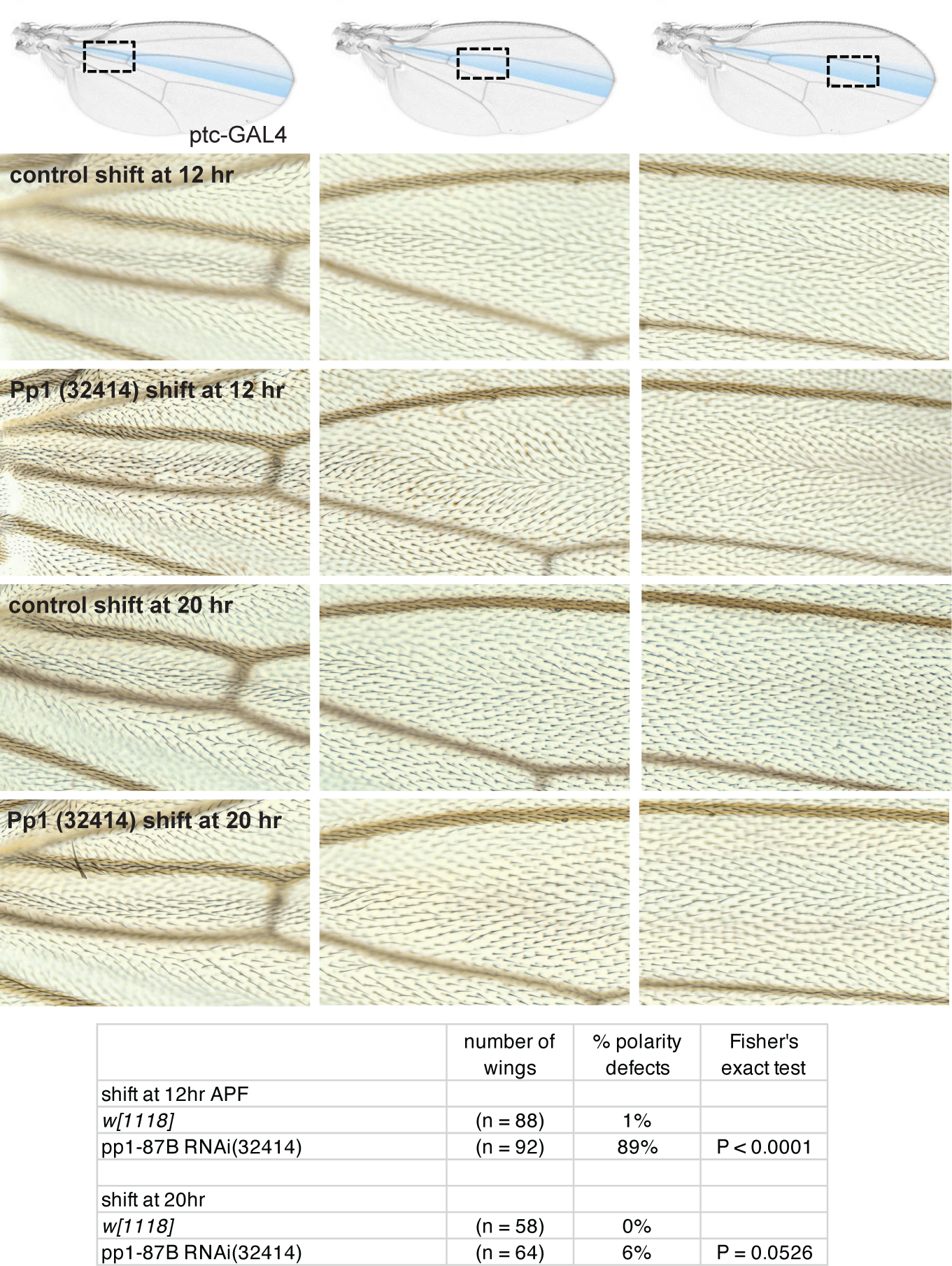
Pp1-87B knockdown in the *ptc* domain disrupts planar polarization. Control (*w^1118^*) or *Pp1-87B* (TRiP RNAi stock 32414) knockdown in the *ptc* domain at either 12 or 20 hours APF. Three regions of the *ptc* domain proximal or distal to the anterior crossvein (acv) are shown. Knockdown by temperature shift to the permissive temperature (18° to 29°) at the equivalent of 12 hr APF but not at 20 hr APF disrupted normal distal polarity, more prominently proximally than distally. The table indicates the percentage of wings that show a polarity perturbation. Scale bar = 50μm.

### Pp1 regulates polarization and localization of Vang

Proper subcellular localization of core PCP proteins correlates with, and is almost certainly responsible for, their ability to direct polarization. If Pp1-87B affects wing hair polarity through the core PCP signaling machinery, it would be expected to alter the characteristic asymmetric subcellular localization of the core proteins to proximal or distal sides of the cell. To examine this, we again used temperature shifts (18° to 29°) to knock down Pp1-87B activity in the *ptc* domain of pupal wings expressing Vang::YFP beginning at 0 hr or 20 hr APF, and harvested at the equivalent of 28 hr APF. Vang::YFP within the *ptc* domain appeared less polarized than a control region just posterior to the fourth vein (**Figure 2**). To quantify this result, the Fiji plug-in ‘TissueMiner’ (Aigouy *et al*, 2010; Etournay *et al*, 2016) was used to define and extract nematics where the length represents the degree of polarization and the angle represents the orientation of polarization in each cell. Using a custom MATLAB script (see Methods) to extract the lengths and compare means, we found that the degree of polarization in the *ptc* domain where Pp1-87B activity is reduced is significantly less compared to the control region outside the *ptc* domain when knockdown is initiated at 0 hr but not at 20 hr APF. We similarly compared orientation of polarization by comparing angle distributions and found a significantly more dispersed orientation distribution in the Pp1-87B knockdown domain compared to the control domain when knockdown is initiated at 0 hr but not at 20 hr APF. Pp1-87B activity is therefore required before 20 hr APF for complete polarization of Vang, and by proxy, the other core PCP signaling proteins.

**Figure 2.**
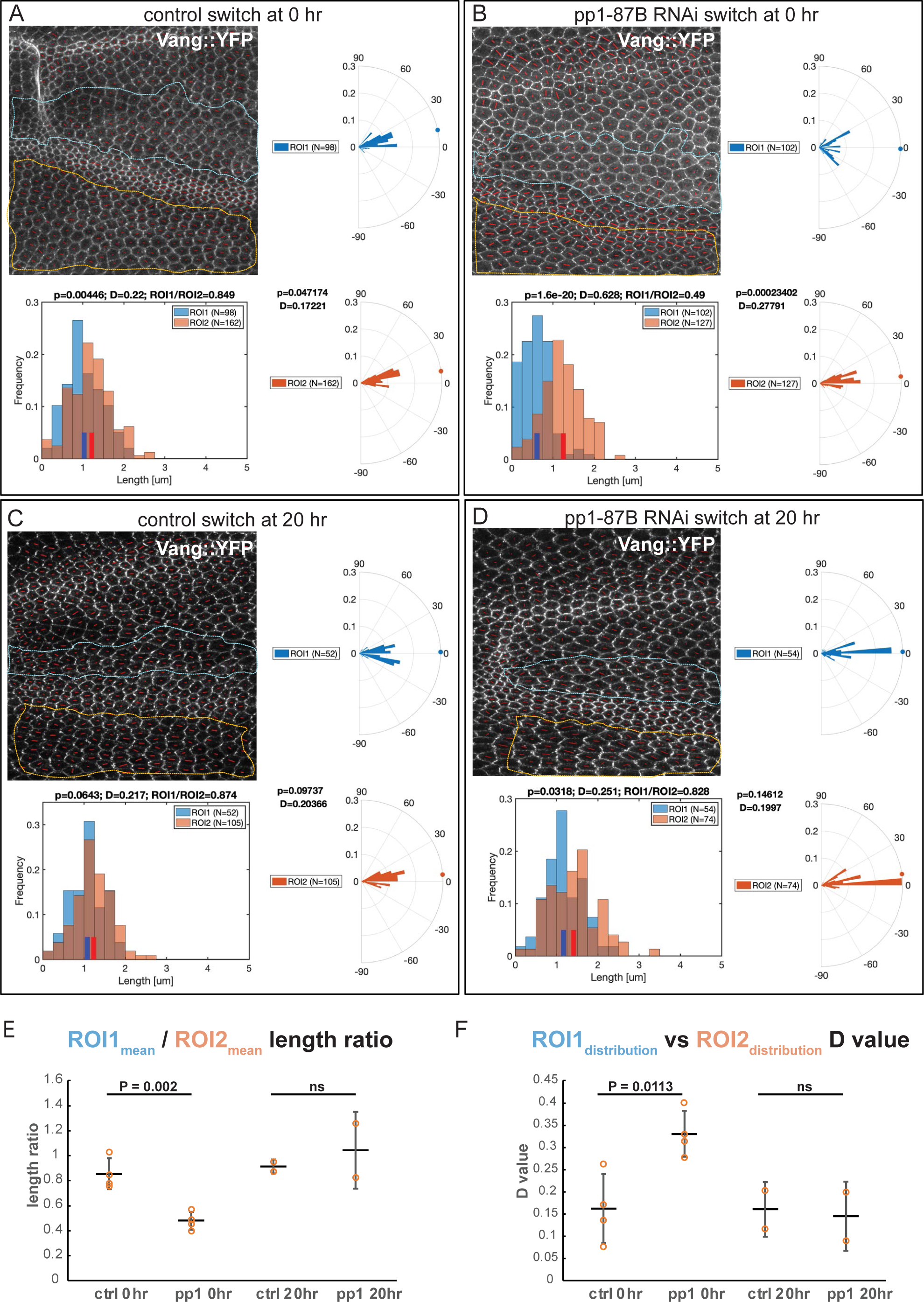
Early but not late knockdown of *Pp1-87B* in the *ptc* domain disrupts polarization of Vang subcellular localization. Knockdown (TRiP RNAi stock 32414) by temperature shift (18° to 29°) was effected in wings expressing Vang::EYFP at the equivalent of 0 or 20 hr APF and pupal wings were harvested at the equivalent of 28 hr APF. The degree of polarization and orientation of polarization of Vang::EYFP was compared for cells in a portion of the *ptc* domain, defined by mRFP expression (ROI1), and an adjacent domain posterior to the L4 vein where no knockdown occurred (ROI2). Polarization of individual cells was determined using the the Fiji plug-in ‘TissueMiner’ (Aigouy *et al*., 2010; Etournay *et al*., 2016) to determine nematics (red overlay lines), and both the length and orientation of each nematic was extracted using a custom MATLAB script (see Methods). Nematic lengths are plotted in histograms and orientations in rose plots. Individual examples are shown in **A-D**. N indicates the number of cells included in the region of interest (ROI). **E**. Ratios of mean nematic length, indicating degree of polarization in the *ptc* domain compared to the posterior domain, for four wings each shifted at 0 hr APF and two each at 20 hr APF are plotted. Shift at 0 hr but not 20 hr produced a significant decrease in mean length ratio (degree of polarization) (P = 0.002, Students t-test). **F**. Distributions of polarity orientation for the *ptc* domain and the posterior domain were compared for the same four wings each shifted at 0 hr APF and two each at 20 hr APF using the Kolmogorov-Smirnov test where D close to 0 indicates identical and D closer to 1 indicates different distributions. D values for shift at 0 hr but not 20 hr APF are significantly greater (P = 0.011, Students t-test), confirming the visual impression that orientations are more dispersed in the *ptc* domain compared to the posterior domain. Scale bar = 20μm.

In similar wings (18° to 30° at 0 hr APF), we also noticed that apicolateral junctional accumulation of Vang::YFP was increased where Pp1-87B activity was reduced, and excess Vang accumulated along the lateral and basal membranes in these cells as well (**Figure 3**). This does not appear to result from gross changes to cell and tissue architecture. A similar result is observed in third instar wing discs shifted to the permissive temperature 48 hr prior to examination (**Expanded View Figure EV4**). We examined the other core PCP proteins for subcellular localization in pupal wings and found that Pk, like Vang, accumulates apicolaterally and laterally, and is also at higher levels throughout the cytoplasm. However, Fz, Dsh and Fmi apical-basal localizations are minimally affected, if at all. We conclude that in addition to perturbing polarization, Pp1-87B knockdown regulates the quantity of the proximal proteins Vang and Pk that can accumulate both apicolaterally and basally.

**Figure 3.**
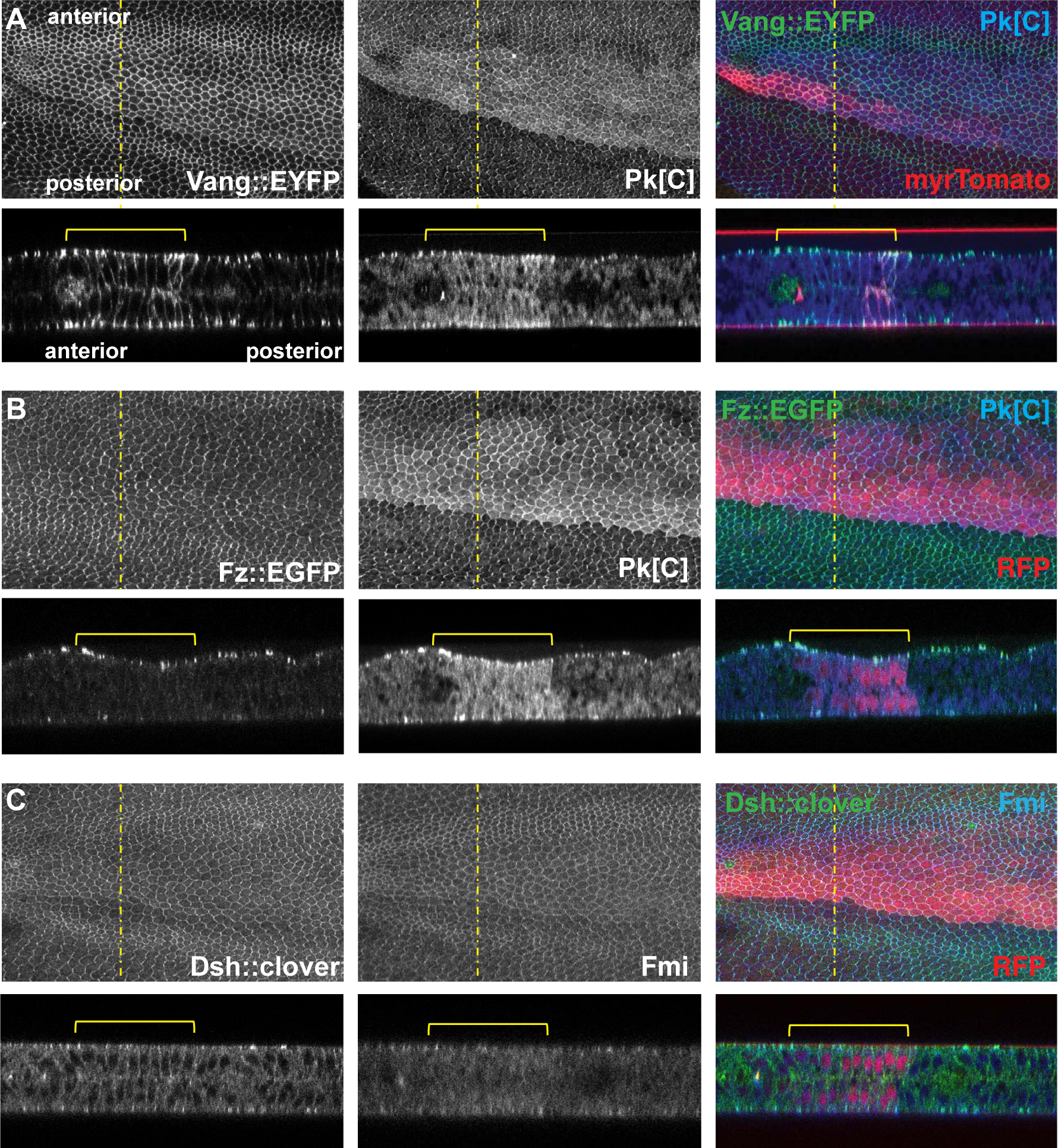
Excess junctional Vang and cytoplasmic Pk accumulation upon *Pp1-87B* knockdown. Pupal wings with *Pp1-87B* knockdown in the *ptc* domain (TRiP RNAi stock 32414) by temperature shift (18° to 30° at 0 hr APF) as in Figure 2, were examined for Vang::EYFP (**A**), Pk (**A** and **B**) (antibody staining), Fz::EGFP (**B**), Dsh::clover (**C**) and Fmi (**C**) (antibody staining) in the X-Y and Y-Z planes. Excess Vang and Pk are observed in apicolateral junctions and along lateral membranes where *Pp1-87B* is knocked down, whereas Fz, Dsh and Fmi are not appreciably affected. The *ptc* domain is marked by myr-Tomato or RFP as indicated. At least 10 wings of each genotype were examined. Scale bar = 50μm.

### Pp1-87B genetically interacts and colocalizes with core PCP components

If Pp1-87B regulates the function of one or more core PCP proteins, either directly or indirectly, it might interact genetically with them. We produced a weak Pp1-87B RNAi phenotype in the *ptc* domain by shifting flies from the restrictive to a partially permissive temperature (18° to 25°) at 0 hr APF. We found that this phenotype was substantially enhanced by heterozygosity for *vang*, though not for *fz* (**Figure 4**). While the negative result for *fz* is not informative, enhancement by *vang/+* provides additional evidence that Pp1-87B regulates core PCP function. Consistent with this possibility, HA-tagged Pp1-87B expressed in the *ptc* domain is seen at apicolateral cell junctions where core PCP proteins localize as well as throughout the cytoplasm (**Appendix Figure S3**). Expression of Pp1-87B in the *ptc* domain subtly perturbed polarity, if at all, and it did not significantly modify Vang polarization or apical-basal localization (**Appendix Figure S4**).

**Figure 4.**
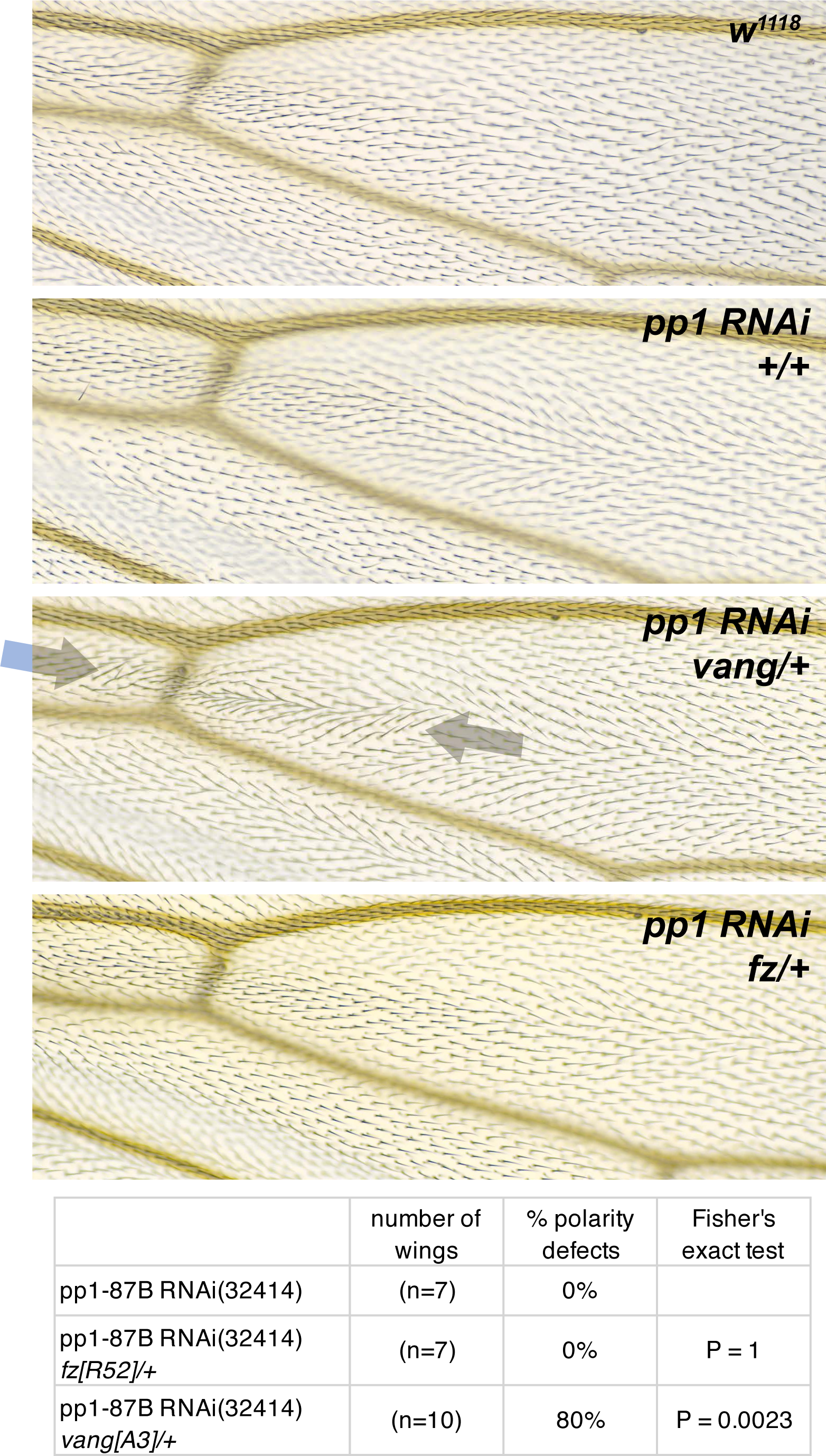
*Pp1-87B* knockdown genetically interacts with *vang*. A weak *Pp1-87B* RNAi knockdown phenotype in the *ptc* domain was produced by temperature shift (18° to 25°) at 0 hr APF. The aberrant polarity was enhanced by heterozygosity for *vang^A3^* but not *fz^R52^*. For consistency, the analysis was limited to the dorsal side of the wing. The table indicates the percentage of wings that show a polarity perturbation. Scale bar = 50μm.

### Pp1-87B genetically interacts with Dco kinase

A number of serine/threonine kinases have been associated with PCP signaling, including PAR- 1, aPKC, CKI: /Dco, CK1δ, CK1ψ/Gilgamesh, Ror2, Nek, Nemo, Drok and Mink1(Collu *et al*, 2018; Daulat *et al*, 2012; Djiane Yogev & Mlodzik, 2005; Gao *et al*, 2011; Gault *et al*, 2012; Kelly *et al*, 2016; Klein *et al*, 2006; Ossipova *et al*, 2005; Schertel *et al*, 2013; Strutt Gamage & Strutt, 2019; Strutt Price & Strutt, 2006; Weber & Mlodzik, 2017; Winter *et al*, 2001; Yang *et al*, 2017). Dco dependent phosphorylation of both the core proteins Dsh and Vang has been demonstrated, although evidence that their phosphorylation by Dco is direct is not definitive. We therefore asked if Dco might interact with Pp1-87B to regulate PCP signaling. Knockdown of Pp1-87B in the *ptc* domain (18° to 29°) at 12 hr APF induced a strong polarity phenotype in adult wings (**Figure 1**), while knockdown at 0 hr APF (18° to 31°) did not yield viable adults. Dco knockdown in the *ptc* domain under these conditions did not perturb polarity, but when we knocked down Dco together with Pp1-87B (0 hr APF, 18° to 31°), the lethality induced by Pp1- 87B knockdown was strongly suppressed and the adult wing hairs showed only a subtle polarity defect (**Figure 5A**). Furthermore, simultaneous Dco knockdown partially suppressed the apical-basal mislocalization of Vang (**Figure 5B,C**). To rule out the possibility that Dco suppressed the Pp1-87B lethality and polarity phenotype by modulating *ptc* expression, we did a similar experiment substituting actin-GAL4 for ptc-GAL4 and saw a similar result (**Expanded Vied Figure EV3**). Together, these results suggest that Dco and Pp1-87B intersect in their regulation of PCP.

**Figure 5.**
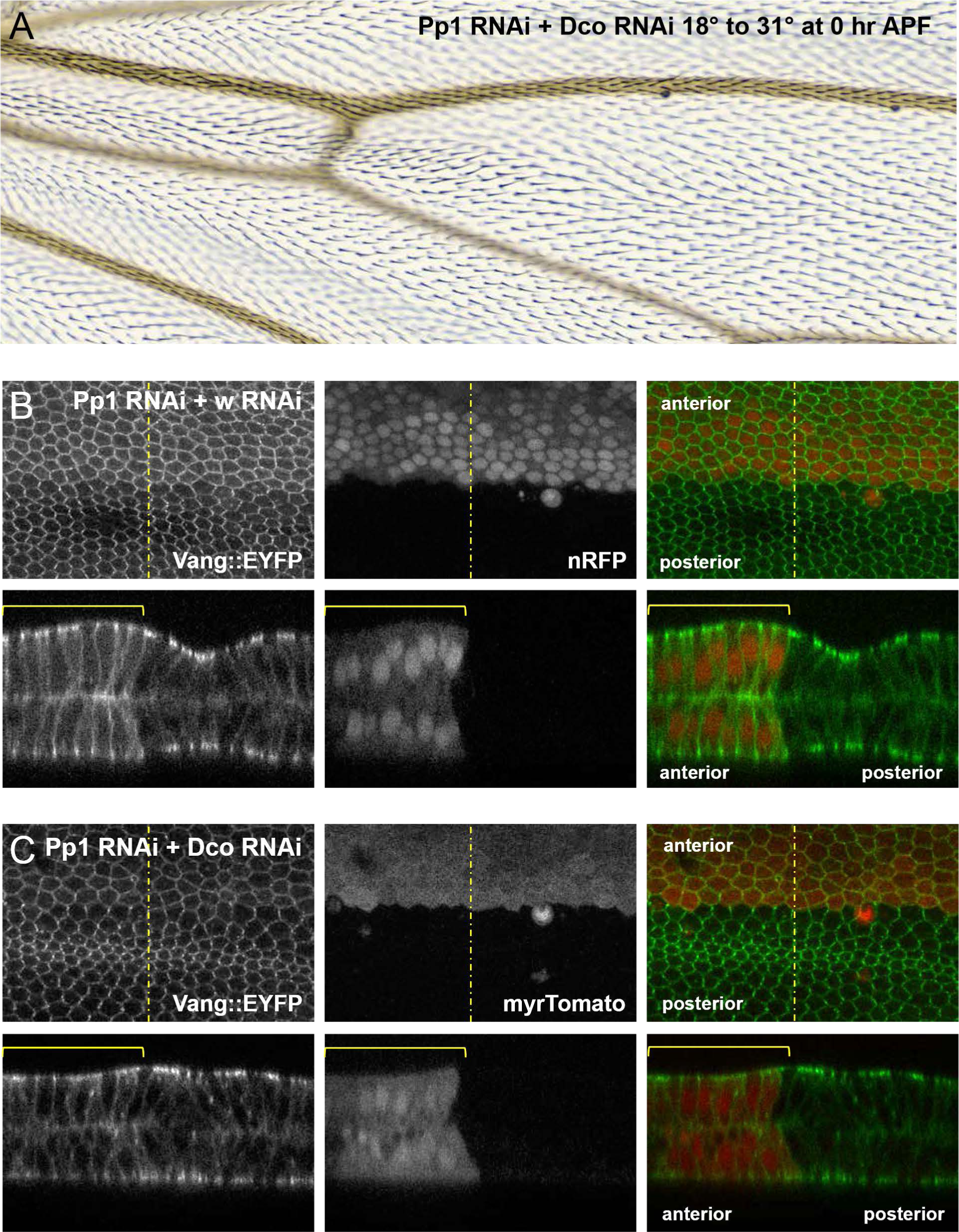
*dco* knockdown suppresses the *Pp1-87B* knockdown phenotype. **A**. *Pp1-87B* RNAi (TRiP RNAi stock 32414 plus wRNAi) knockdown at 0 hr APF produces lethality, but knockdown of Dco together with knockdown of Pp1-87B (TRiP RNAi stock 32414 plus *dco* RNAi) rescues viability and produces only a weak PCP phenotype. **B-C**. Vang::EYFP accumulates laterally and basally in 28 hr APF pupal wings when *Pp1-87B* RNAi is knocked down together with white (wRNAi) at 0 hr APF as in (A); co-expression of *wRNAi* somewhat weakens the effect compared to *Pp1-87B* RNAi alone (Figure 1), likely due to GAL4 dilution. Knockdown of dco (dcoRNAi) together with *Pp1-87B* RNAi partially suppresses lateral and basal Vang::EYFP accumulation relative to the control. At least 10 wings of each genotype were examined. Scale bars = 50μm.

### Pp1-87B modifies phosphorylation state of Vang in vivo

Our results thus far indicate that Pp1-87B regulates core PCP polarization (**Figure 2**) and genetically interacts with *vang* but not *fz* (**Figure 4**), and with Dco, a kinase thought to phosphorylate Vang (**Figure 5**). Furthermore, the subcellular localization of Vang is modified by knockdown of Pp1-87B (**Figure 3**). We therefore sought to determine whether Pp1-87B might modify the phosphorylation status of Vang in vivo. Extracts from third instar wing discs expressing Pp1-87B RNAi (32414 or 67911) or overexpressing Pp1-87B, each under the control of 71B-GAL4 were probed by Western blot (**Figure 6**). Control extract treated with lambda protein phosphatase showed a modest shift toward faster mobility, consistent with dephosphorylation of Vang. We detected a similar though perhaps more subtle shift in the Pp1- 87B overexpression condition. We could not reliably detect a shift toward hyperphosphorylation upon Pp1-87B knockdown. Similarly, no shift was detectable with Dco knockdown, as has been reported by others, although such a shift was detected in transfection experiments (Kelly *et al*., 2016). These results are consistent with Vang being largely in a phosphorylated state in vivo, and with Vang being a substrate for Pp1-87B phosphatase activity in vivo, although it is also possible that Pp1-87B indirectly dephosphorylates Vang.

**Figure 6.**
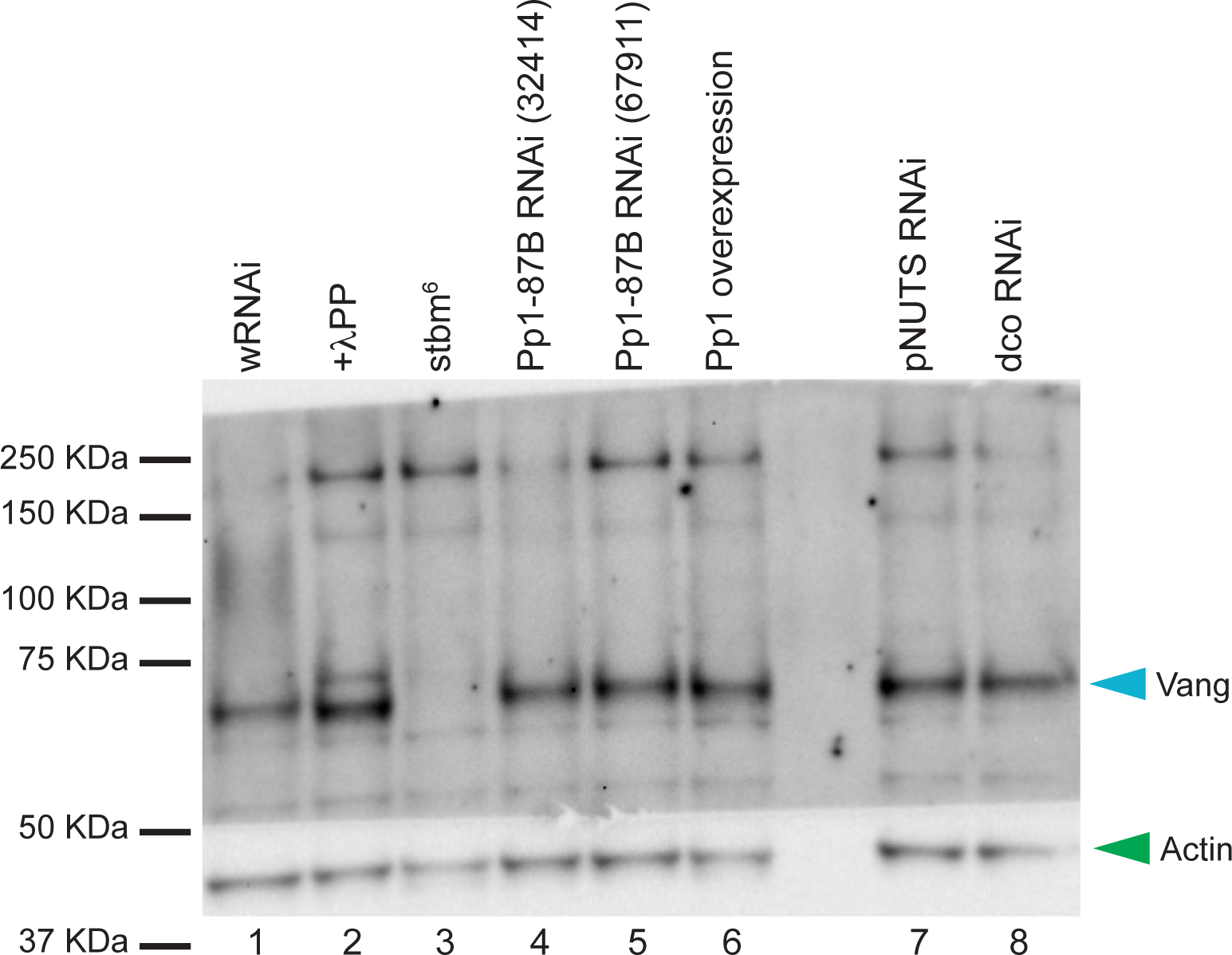
Pp1-87B overexpression reduces phosphorylation of Vang. Vang runs as a single band (lane 1), and with 11 Protein Phosphatase treatment, runs faster, showing that it is phosphorylated in vivo (lane 2). We were unable to detect a shift toward hyperphosphorylation of Vang from wing discs expressing either of two Pp1-87B RNAi lines throughout the wing under 71B-GAL4 control (lanes 4 and 5), but Vang from wing discs broadly overexpressing Pp1::HA, Vang migrates slightly faster, suggesting less phosphorylation. We did not detect a shift from wing discs expressing pNUTS or dco RNAi. All extracts are from third instar wing discs. Actin was probed as a loading control. Two biological replicates were performed for this experiment.

### PNUTS is a potential subunit of the Pp1 holoenzyme in PCP signaling

Pp1 functions as a holoenzyme, deriving substrate specificity by partnering with specific non-catalytic subunits to find and act on its substrates in various contexts(Bertolotti, 2018; Peti Nairn & Page, 2013). In vertebrates, known non-catalytic subunits comprise a large family of structurally diverse proteins; in *Drosophila*, a modest number of non-catalytic subunits has been identified(Bennett *et al*, 2006). Several known subunits were identified in our mass spec results, though they scored below our significance threshold: we were unable to produce a convincing polarity phenotype upon knockdown of Ankyrin-repeat, SH3-domain, and Proline-rich-region containing Protein (ASPP), Myosin binding subunit (Mbs), or Mars with any available RNAi line or driver we tested. Knockdown of Nebbish (Neb) produced a weak multiple wing hair phenotype but at most a marginal hair orientation defect. We then examined several other known Pp1-87B interacting proteins. Knockdown of Phosphatase 1 nuclear targeting subunit (PNUTS)(Ciurciu *et al*, 2013) in the *ptc* domain induced a clear polarity phenotype proximal to the anterior crossvein (acv), where hypomorphic dco allelic combinations and two originally identified antimorphic Vang alleles have been shown to disrupt polarity (Strutt Price & Strutt, 2006; Taylor *et al*., 1998) (**Figure 7 and 8**). This proximal phenotype differed in pattern on the dorsal and ventral sides of the wing, and notably, neither disruption extended substantially distal to the acv, where we see a robust Pp1-87B knockdown phenotype. We were, however, able to detect a modest disruption to the polarity nematic distal to the acv (**Expanded View Figure EV5**). This was not accompanied by a detectable change in apical or basal Vang::GFP accumulation (**Appendix Figure S4**).

**Figure 7.**
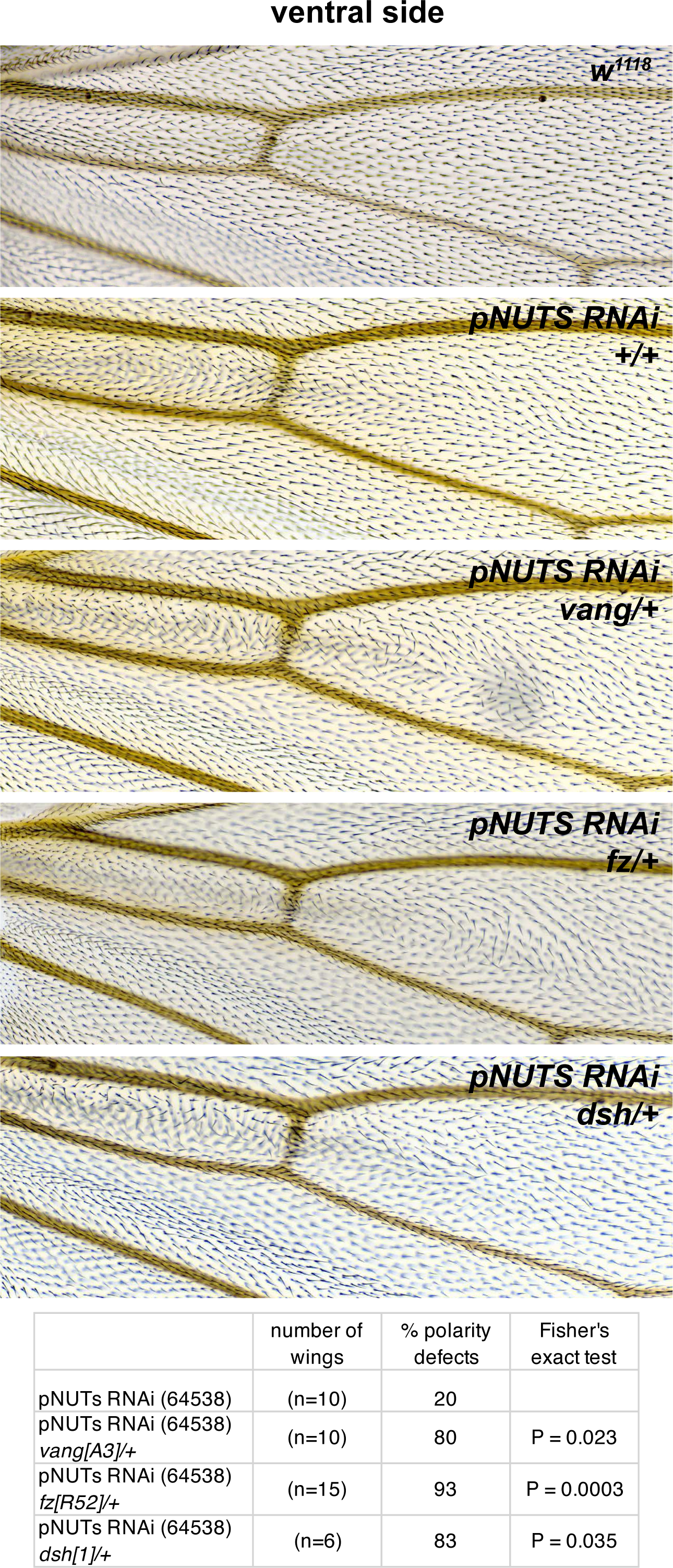
*PNUTS* knockdown interacts with core PCP genes *vang, fz* and *dsh*. Because *PNUTS* knockdown (TRiP RNAi stock 64538) in the *ptc* domain produces substantially different patterns on the dorsal and ventral sides, we restricted interaction analyses to a single side, in this case ventral. The weaker ventral phenotype is enhanced by heterozygosity for *vang^A3^*, *fz^R52^* and *dsh^1^*. The table indicates the percentage of wings that show a polarity perturbation. Scale bar = 50μm.

**Figure 8.**
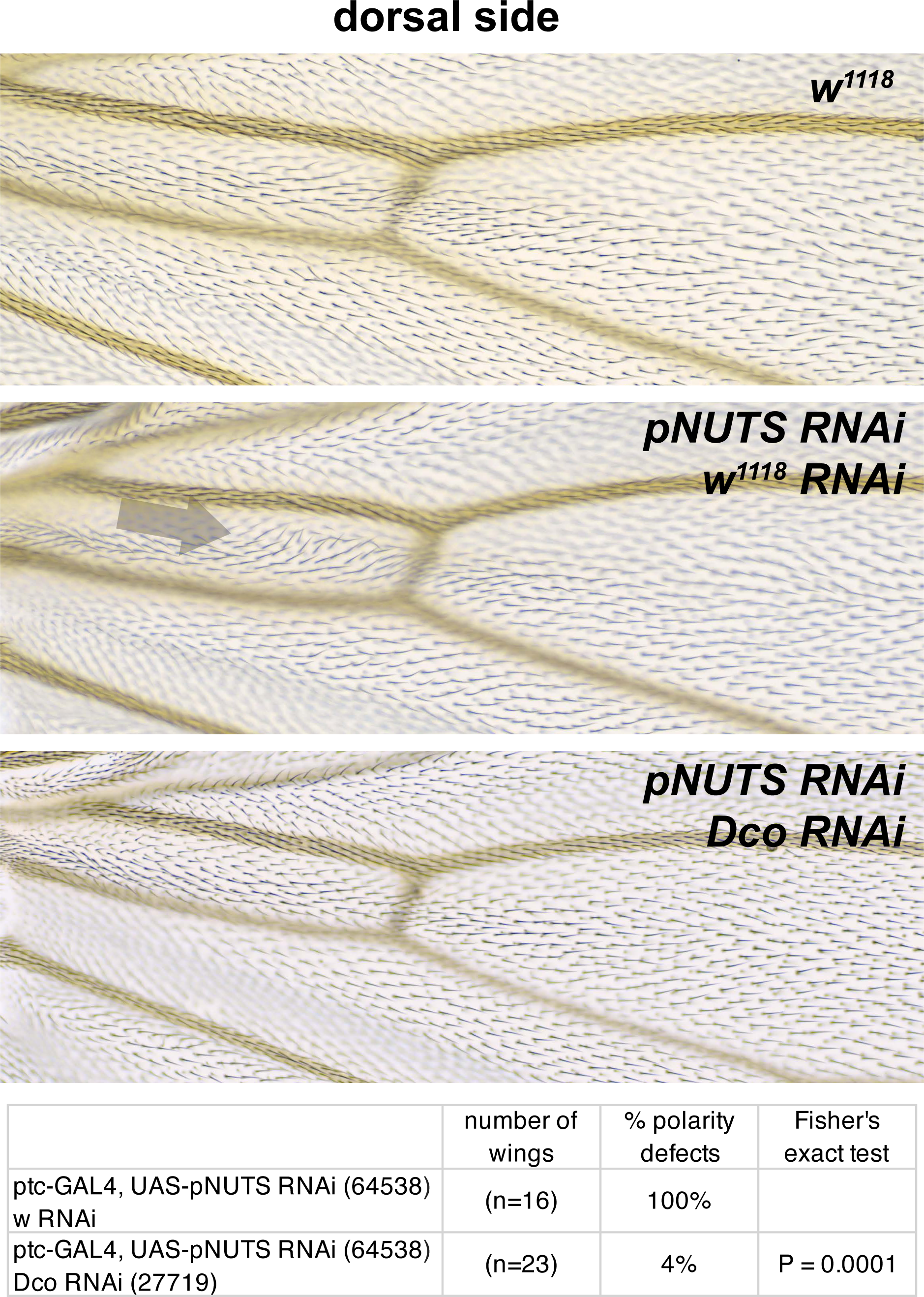
*PNUTS* knockdown interacts with *dco* knockdown. The dorsal *PNUTS* knockdown (TRiP RNAi stock 64538) phenotype in the *ptc* domain (arrow) was suppressed by simultaneous *dco* knockdown. Because *PNUTS* knockdown in the *ptc* domain produces substantially different patterns on the dorsal and ventral sides, we restricted interaction analyses to a single side, in this case dorsal. The table indicates the percentage of wings that show a polarity perturbation. Scale bar = 50μm.

The PNUTS polarity phenotype was enhanced by heterozygosity of *vang, fz* and to a lesser extent, *dsh*, and the enhanced polarity disruption in many instances extended distal to the position of the acv (**Figure 7**). Like Pp1-87B, the PNUTS knockdown phenotype was suppressed by simultaneous Dco knockdown (**Figure 8**) Due to the different dorsal and ventral polarity patterns on PNUTS inhibition, we restricted each of these interaction analyses to a single side. No shift in Vang phosphorylation was detected on pNUTS knockdown, though this was not expected given no shift could be detected with Pp1-87B knockdown (**Figure 6**) Thus, reduction of PNUTS results in many, though not all, of the alterations seen with reduction of Pp1-87B, except that they are shifted toward a more proximal position in the wing.

## Discussion

While the dedicated components of the core PCP mechanism have been known for some time, regulators that facilitate their activity, and which are likely to have pleiotropic effects, have been more challenging to identify in genetic screens. We employed an APEX based proximity labeling approach coupled with iTRAQ and LC-MS/MS to identify such regulators. Tagging Vang with APEX led to the identification of 123 high probability hits; these, plus selected lower probability hits, were secondarily screened by RNAi, and 91 showed either mis-orientation of wing hair polarity or multiple wing hair phenotypes (or both). The true positive rate may be higher given the variable efficiency of RNAi lines. Our screen therefore provides a resource for exploring previously uncharacterized regulators of PCP signaling.

Among the positive hits, we chose to further characterize the role of the S/T phosphatase catalytic subunit Pp1-87B. Previously, Widerborst (wdb), a regulatory subunit of PP2A was identified as a regulator of PCP, though its precise role is poorly defined (Hannus *et al*, 2002).

Numerous post-translational modifications of core PCP components have been proposed to control core PCP signaling, including phosphorylation of *Drosophila* and vertebrate homologs of Fmi, Fz, Vang, Pk and Dsh (reviewed in (Harrison *et al*., 2020)). Of these, phosphorylation of Dsh and Vang has been most thoroughly investigated.

A phosphorylation dependent gel shift of Dsh is associated with PCP signaling (Axelrod, 2001; Shimada *et al*, 2001). CKI: /Dco, a S/T kinase, is required for PCP signaling in flies and vertebrates, and CKI: has been found to bind to and phosphorylate vertebrate Dishevelled proteins (Dvl) in numerous cell culture and in vitro assays (Cong Schweizer & Varmus, 2004; Gao *et al*, 2002; Hino *et al*, 2003; Kishida *et al*, 2001; McKay Peters & Graff, 2001; Peter *et al*, 1990; Sakanaka *et al*, 1999). *Drosophila* Dco is similarly able to phosphorylate Dsh in vitro, and mutation of S236, in a consensus motif predicted to be phosphorylated by Dco, blocks that ability (Klein *et al*., 2006; Strutt Price & Strutt, 2006). Surprisingly, however, in S2 cells a kinase- dead CKI: induces Dsh phosphorylation as efficiently as does the functional kinase, suggesting that Dco, via a kinase-independent mechanism, stimulates one or more other kinases to phosphorylate Dsh (Klein *et al*., 2006). In a cell culture model, MS mapping of Dsh phosphorylation identified many Fz-dependent and independent S/T phosphorylations (though surprisingly, not S236) (Yanfeng *et al*, 2011). However, none of these, either alone or in several combinations, were shown to be functionally important for PCP signaling in an in vivo rescue assay, although substantial redundancy cannot be ruled out. Furthermore, mutation of the of S/T target cluster in the C-terminus did not disrupt rescue activity, and indeed the C-terminus of Dsh, which accounts for the bulk of the phosphorylation-dependent gel shift, is not required for rescue of PCP signaling. Thus, despite the extensive S/T phosphorylation of Dsh, these attempts failed to define functional significance for Dsh phosphorylation. In contrast, two subsequent observations provide at least weak evidence for functional significance: a modest genetic interaction between overexpressed Dco and *dsh* was observed, and a phosphomutant including S236 and seven adjacent S/T residues that failed to significantly disrupt polarity nonetheless showed a subtle decrease in Fmi asymmetry (Strutt Gamage & Strutt, 2019; Strutt Price & Strutt, 2006). In sum, while Dsh S/T phosphorylation correlates with membrane localization, functional significance of this phosphorylation remains equivocal.

Of note, the MS analysis identified a tyrosine phosphorylation (Y473) that was shown to be mediated by the Abl kinase and to have a relatively PCP-specific requirement, as a Y473F mutant behaves much like the PCP-specific *dsh^1^* allele (Singh *et al*, 2010; Yanfeng *et al*., 2011). Tyrosine phosphorylations have been proposed to regulate additional aspects of PCP signaling as well (Guo Zanetti & Schekman, 2013; Humphries *et al*, 2023; Kelly *et al*., 2016).

*Drosophila* and vertebrate Vang phosphorylation appears to be mediated by CKI:/Dco, likely directly (Gao *et al*., 2011; Kelly *et al*., 2016; Yang *et al*., 2017). Two clusters of S/T phosphorylation sites were identified in Vang, and their mutation disrupts PCP in flies and vertebrates (Gao *et al*., 2011; Kelly *et al*., 2016; Ossipova Kim & Sokol, 2015; Strutt Gamage & Strutt, 2019; Yang *et al*., 2017). Because mutation to either phosphomutant or phosphomimetic residues disrupts PCP, an appealing explanation is that cycling between phosphorylated and unphosphorylated forms may be required (Strutt Gamage & Strutt, 2019). Based on measurements of stability and localization asymmetry, it was proposed that phosphorylation destabilizes Vang in puncta, and that cycling between phosphorylated and unphosphorylated forms is required for the mutual inhibition of oppositely oriented complexes that generates asymmetry (Butler & Wallingford, 2017; Strutt Gamage & Strutt, 2019). This model is supported by the observation that phosphorylation of Vang is cell autonomously stimulated by Fz, and cell autonomous interaction between Fz and Vang complexes is hypothesized to destabilize one or the other (Amonlirdviman *et al*, 2005; Kelly *et al*., 2016; Tree Ma & Axelrod, 2002). Consistent with this idea, Pk was proposed to stabilize Vang by inhibiting its phosphorylation (Strutt Gamage & Strutt, 2019), but alternately, or perhaps in addition, Pk was proposed to destabilize Vang by promoting its internalization (Cho *et al*, 2015).

While Dco may participate in cycling Vang between phosphorylated and less phosphorylated states to regulate PCP, a corresponding phosphatase activity has not been reported. Here, we’ve provided evidence that the S/T phosphatase catalytic subunit Pp1-87B might participate in such a role. Appropriately timed reduction of Pp1-87B expression produces a polarity phenotype consistent with this function as assessed by wing hair polarity and subcellular localization of core PCP proteins, and by genetic interaction with Vang. Antagonistic interaction between Pp1-87B and Dco to modulate PCP further suggests that this kinase and Pp1-87B oppositely regulate a common process. Vang accumulates and is mislocalized upon Pp1-87B knockdown, consistent with its localization in puncta being destabilized by Fz-dependent phosphorylation and stabilized by dephosphorylation.

Biochemical evidence for these activities is incomplete. Prior studies of Vang phosphorylation showed that upon co-transfection into S2 cells with Dco, but not a kinase dead Dco, a shift reflecting presumptive phosphorylation was detected (Kelly *et al*., 2016). However, larval lysates from *dco^2^* or *dco^3^* hypomorphic mutants (Kelly *et al*., 2016) or Dco knockdown (**Figure 6**) did not show an appreciable shift. This may be due to remaining activity of the hypomorphic alleles or incomplete knockdown, or redundant activity of another kinase in vivo. It is also likely that the contribution of Dco-dependent phosphorylation is a modest fraction of the total phosphorylation of Vang in vivo, making it difficult to detect. Treatment with the dual serine/threonine and tyrosine specific lambda phosphatase reveals significant phosphorylation of Vang in vivo, and tyrosine phosphorylation of Vang has also been demonstrated (Humphries *et al*., 2023). The subtle shift in phosphorylation we detect with Pp1-87B overexpression, and lack of shift upon knockdown, are consistent with the possibility that Pp1-87B phosphatase activity, in conjunction with pertinent regulatory components, acts on a limited set of Vang phosphorylation sites in the context of PCP signaling in vivo. Interestingly, *pk* mutants show a shift in Vang toward hyperphosphorylation, suggesting that Pk antagonizes Vang phosphorylation (Strutt Gamage & Strutt, 2019), and the magnitude of this effect raises the possibility that it does more that impact phosphorylation of the sites potentially regulated by Dco and Pp1-87B.

In sum, our data suggest that Pp1-87B regulates the phosphorylation state of Vang, though we have not determined whether this action is direct or indirect. As Pp1-87B is thought to be the most abundant and active S/T phosphatase (Dombradi *et al*., 1990), it would be unsurprising if it has multiple targets in the core PCP pathway. Cycling of Vang between phosphorylated and dephosphorylated states to facilitate mutual exclusion is expected to be contextually regulated: appropriately localized activity of Dco and the corresponding phosphatase Pp1 may fulfill this function.

That Vang accumulates and is mislocalized upon reduction of Pp1 activity is not simply predicted or explained by current understanding. Reduction of phosphorylated Vang’s residence time in puncta with oppositely oriented complexes (Strutt Gamage & Strutt, 2019) does not necessarily imply that the protein is itself destabilized, and indeed if excess Vang is expelled from such puncta, it may accumulate ectopically in cells. Similarly, excess phosphorylated Vang, perhaps having diminished residency in large puncta, might exit those large puncta and seed the formation of new, small puncta at apicolateral junctions. Furthermore, action of Pp1 on substrates other than Vang or its regulators might contribute to the phenomenon. The fate of this phosphorylated Vang population will require further study.

The substrate specificity of protein phosphatases is controlled by non-catalytic subunits. While the catalytic subunits of Pp1 alone are highly promiscuous, in cells they exist almost exclusively as subunits of a holoenzyme in which other subunits provide a high degree of substrate specificity (Bertolotti, 2018; Brandt Killilea & Lee, 1974). Hundreds of human regulatory subunits exist, and at least 40 putative regulatory subunits of perhaps many more in *Drosophila* have been identified (Bennett *et al*., 2006; Bertolotti, 2018; Hendrickx *et al*, 2009). Though no known Pp1 regulatory subunit reached significance in our screen, we identified a PCP phenotype with PNUTS knockdown, and interactions between PNUTS and other PCP components are quite similar to those of Pp1. Given the more limited territory in the wing in which PNUTS appears to act, contributions of other regulatory subunits are likely also be involved.

Our data suggest a role for PNUTS in regulating PCP. Since the most dramatic effects of Pp1 are observed on Vang, one might speculate that PNUTS targets Pp1 to Vang. However, unlike Pp1, PNUTS knockdown does not regulate Vang accumulation or apical basal localization.

Furthermore, PNUTS was previously shown to target Pp1 activity to RNA polymerase II (Ciurciu *et al*., 2013), raising the alternate possibility that PNUTS indirectly regulates PCP through broader changes in transcription. Furthermore, ribosome composition dependent translation of Vangl2 and Fz3 in differentiating human embryonic stem cells has recently been recognized, suggesting yet another potential target point (Genuth *et al*, 2022). Additional studies are required to determine how Pp1, PNUTS and/or one or more other regulatory subunits are involved in regulating the core PCP machinery.

## Methods

### APEX::Vang flies

Vang cDNA from fly larvae was cloned into a Gateway pENTR vector and sequence verified. The Vang cDNA was recombined into a pUAST Gateway destination vector with 3xMyc-APEX fused to the N-terminus of Vang. Transgenic flies with site-specific insertion of UAS-3xMyc- APEX-Vang were made by injection of the plasmid together with PhiC31 integrase and targeted for insertion into the attP40 site on the 2^nd^ chromosome.

### APEX labeling and sample preparation for Western blot and proteomic analyses

3^rd^ instar wing discs were biotinylated and prepared in batches for western blots and mass spectrometry as follows. Dissected wing discs were collected into cold labeling buffer (PBS + 500mM biotin-phenol +2mM probenecid) on ice. Aliquots were then transferred to room temp labeling buffer for 30 min. Labeling buffer was removed and 1mM hydrogen peroxide in PBS was added to activate biotinylation for 1 min. The reaction was quenched immediately by adding quenching medium to a final concentration of 10mM sodium azide, 10mM sodium ascorbate and 5mM Trolox in PBS. The tissue was washed in quench medium x3 then PBS x2. The discs were removed into cold PBS with protease inhibitors (Roche) and transferred to a 0.6mL Eppendorf tube. An equal volume of 2x RIPA was added, tissue was hand ground, and then spun at 14,000rpm for 10 min at 4° C. Supernatant was transferred to a new tube, immersed into liquid nitrogen and stored at -80° C. Protein concentration was determined using a BCA kit (Pierce).

### Streptavidin bead binding

Streptavidin magnetic beads (Pierce; amount with sufficient binding capacity for protein amount) were washed with PBS, then with RIPA. Protein in RIPA was added to the washed beads and allowed to bind for 1 hour at room temperature. Beads were then washed in RIPA 3 times. For Western blotting, 60μL of 1x SDS page buffer was added to the beads and heated for 5min at 95° C. The elution buffer E1, was removed from the beads.

### Western blotting for Fmi and Pk

Samples were loaded on a Tris acetate 3-8% gel beside HiMark prestained protein standard was also loaded. The gel was run for 2 hours and then proteins were transferred to a PVDF membrane. The membrane was blocked for one hour in either, 5% dry milk in 50mM Tris, 150mM sodium chloride (blocking buffer) for Fmi, Myc and Pk, or in Pierce protein free block for streptavidin. Membranes for Fmi, Myc and Pk were probed with antibodies overnight at 4° C or with streptavidin as below for 1 hour. Membranes were then washed, ECL applied and then exposed to x-ray film.

Fmi and Myc:

20μL (13μg) of input was loaded for each genotype/lane and the same volume of E1 (20μL) was also loaded.

After transfer, the membrane was cut horizontally at 117kDa.

The upper portion was incubated with Fmi antibody and the lower portion with Myc antibody. Primary: anti Flamingo #74-c (DSHB) 1:40 in 5% blocking buffer Secondary: donkey anti mouse HRP 1:5,000 in 5% blocking buffer Primary: anti Myc (source) 1: 5,000 in 5% blocking buffer Secondary: donkey anti rabbit HRP 1:5,000 in 5% blocking buffer

For Fmi, Western Bright ECL K12045-D20 sensitive HRP substrate for chemiluminescent Western blots was used. For myc, Supersignal West Pico chemiluminescent substrate was used.

Pk:

15μL (8μg) of input was loaded for each genotype/lane and the same volume of E1 (15μL) was also loaded.

Primary: guinea pig anti pk (RRID:AB_2941929 (Olofsson *et al*, 2014)) 1:2,000 in 5% blocking buffer

Secondary: donkey anti guinea pig HRP 1:10,000 in 5% blocking buffer

Proteins were visualized with Supersignal West Pico ECL.

Streptavidin:

12μL (1.5μg) of input was loaded for each genotype/lane and the same volume of E1 (12μL) was also loaded.

After blocking for 1 hour with Pierce protein free block, the membrane was probed with streptavidin horse radish peroxidase (SA10001) 1: 50,000 in Pierce block for 1 hour at RT.

Proteins were visualized with Supersignal West Pico ECL.

### Samples for mass spectrometry

2 independent samples were generated for both genotypes, D174gal4 and D174gal4>APEX::myc::vang. Batches of biotinylated discs were prepared as above and numbered sequentially on the day of dissection and labeling. Even numbered and odd numbered tubes were pooled separately for each genotype. The final pooled volumes were approximately 1800μL. The total protein concentration of each sample was measured by BCA. Each sample was then bound to washed streptavidin beads as above. The beads were washed in PBS and sent for mass spec.

### Mass spectrometry

2 mg total input protein of each replicate was enriched for biotinylated proteins using streptavidin beads and analyzed by high-resolution quantitative LC- MS/MS. On-bead digestion was subsequently performed by incubating the beads in trypsin solution (80 µL 1 mM DTT, 5 μg/mL trypsin in 2 M urea in 50 mM Tris, pH 8) overnight to retrieve peptides of biotinylated proteins from the beads for the following analysis. The supernatant was removed and reduced (4 mM DTT), alkylated (10 mM iodoacetamide) for 30 min each, and digested with 0.5 µg of trypsin overnight (37 °C). The resulting digested peptides were desalted and reconstituted in 30 μL iTRAQ reconstitution buffer. Four-plex iTRAQ labeling was conducted per the manufacturer’s instructions (SCIEX). Briefly, iTRAQ labels were reconstituted with ethanol to a final volume of 145 µL, followed by individual labeling at room temperature for 1 h by adding 140 μL iTRAQ reagent to the samples. Labels were used as follows: 114 for APEX::myc::vang, 115 for control, 116 for APEX::myc::vang, and 117 for control replicate B. Label incorporation was evaluated on an Orbitrap before quenching with 100 mM (final) Tris for 10 min at room temperature.

Peptides were separated over a 260-min gradient using a heated PicoFrit (New Objective) column (50C) packed with 24 cm of 1.9-µm C18 material (Dr. Maisch) and run on a Q Exactive MS in data-dependent mode where the top 12 most abundant precursors were selected for MS/MS prior to being placed on an exclusion list. Data were searched with Spectrum Mill (Agilent) using the UniProt *Drosophila* database. A fixed modification of carbamidomethylation of cysteine and variable modifications of N-terminal protein acetylation, oxidation of methionine, and 4-plex iTRAQ labels were searched. The precursor mass tolerance and MS/MS tolerance were set to 20 ppm. The peptide and protein false discovery rates were set to 0.01.

### Proteome analysis and prioritization of hits

Of 2418 proteins identified, 2064 proteins were identified by >1 unique peptide and were selected for further analysis. Identified peptides with UniProt accession numbers were mapped to FlyBase gene identifiers using an ID mapping tool at UniProt that correspond to 2,028 unique annotated *Drosophila* genes (FlyBase release 6.10). All of the genes identified by iTRAQ along with their annotation are listed in Expanded View Dataset 1.

The log2 ratios of each replicate of experimental sample (D174-GAL4, UAS-APEX::Vang; replicates 114 and 116) to each replicate of the control sample (D174-GAL4; replicates 115 and 117) (ratios: 114/115, 116/117, 116/115, and 116/117) were calculated and normalized to the median value of each channel, respectively.

To determine the significance cutoff of the iTRAQ ratio for each dataset, proteins were cross-referenced to positive and negative control lists. The positive control list was assembled manually based on literature, while a list of transcription factors and metabolic genes with high confidence annotation (Hu *et al*, 2015) was used as negative control.

The false positive rate (FPR) is calculated for each iTRAQ ratio using the equation as described in the literature (Chen *et al*., 2015; Rhee *et al*., 2013), where the numerator is the conditional probability of finding a false positive protein in a particular iTRAQ ratio range and is calculated as the percentage of proteins on the negative control list over all proteins identified on negative control list, and the denominator is he percentage of proteins on the positive control list over all proteins identified on the positive control list. The FPR is the ratio of these two conditional probabilities, and is illustrated in the plot in Appendix Figure S5 for each iTRAQ ratio. We selected as our cut-off point for each replicate the iTRAQ ratio at which the FPR reaches 0.1. All proteins above this cut-off are >10 times more likely to be true positives than false positives.

The genes that were above the cutoff in three or four of the four datasets were selected as high confidence hits while genes above the cutoff in 2 datasets were considered as low confidence hits (Dataset 3). Protein complex annotation from COMPLEAT (Vinayagam *et al*, 2013) (www.flyrnai.org/compleat/) was used to identify complexes associated among the genes identified by APEX.

### RNAi screen

The RNAi screen was first performed with constitutive GAL4 drivers (one of the following: *ptc- GAL4, ci-GAL4, hh-GAL4, ap-GAL4; the majority were screened with ptc-GAL4 after initially experimenting with each of them*). Progeny were mostly raised at 29°C; a minority were raised at 25°C as indicated. Those producing lethality were secondarily crossed to *hsFlp>CD2>GAL4* (#133) or *Dsh-clover2; hsFlp>CD2>GAL4* (#476), the 3^rd^ instar larvae were heat-shocked at 37°C for 30 min to induce clones, then raised at 25°C. Adult wings were mounted and examined for polarity phenotypes as described below.

### qPCR protocol

RNA was obtained from 20-30 late third instar wing imaginal discs that were dissected and transferred to ice-cold PBS + 0.2% Triton X-100 (PBS-T). We used *71B-Gal4 / UAS-w RNAi* as control and *71B-Gal4 / UAS-Pp1-87b RNAi (32414)* and *71B-GAL4 / UAS-Pp1-87b RNAi (67911)* to quantify the efficiency of both Pp1 RNAi lines. Discs were spun down at 9000 g for 3 min, after which PBS-T was removed and discs were snap-frozen in liquid nitrogen.

Wing disc total RNA was extracted using the Qiagen RNeasy Mini RNA extraction kit, following manufacturer’s instructions. RNA was then retrotranscribed to cDNA using Thermo Scientific’s SuperScript IV First strand synthesis kit, following the manufacturer’s protocol and using oligo(dT) primers.

The obtained cDNA was then diluted 1/100 and qPCR was performed on an Applied Biosystems StepOne Real-Time PCR machine, using the Applied Biosystems PowerUp SYBR Green master mix. To normalize expression levels, we used RpL32 as reference gene. The following primers were used for the PCR reactions:

RpL32 F: ATGCTAAGCTGTCGCACAAATG

RpL32 R: GTTCGATCCGTAACCGATGT

Pp1 F-I: GATCCGGGGACTTTGCTTGAA

Pp1 R-I: GAGGAACAGGTAATTCGATTCCG

Pp1 F-II: GCAATCGCTGGAGACGATCT

Pp1 R-II: TCAAATCGGGACTGAGACCAC

### Temperature shift protocols

Expression levels of GAL4 driven RNAi lines and other transgenes were controlled by raising flies at defined temperatures through defined developmental time periods. Specific shifts were performed as indicated for each experiment. Because progression through development is temperature dependent, for simplicity, all developmental times are reported as adjusted to the equivalent times at 25°.

### Adult wing hair imaging

Adult wings were dissected and washed with 70% EtOH and mounted in DPX (Sigma) solution. All adult wings were imaged on a Nikon Eclipse E1000M equipped with a Spot Flex camera (Model15.2 64 MP).

### Immunohistochemistry

Pupal wings were dissected at indicated developmental time points after puparium formation (APF). Pupae were removed from their pupal cases and fixed for 60–90 min in 4% paraformaldehyde in PBS at 4°C. Wings were then dissected and extracted from the cuticle, and washed two times in PBST (PBS with 0.1% Triton X-100). After blocking for 1 hr in 5% Bovine serum Albumin in PBST at 4°C, wings were incubated with primary antibodies overnight at 4°C in the blocking solution. Incubations with secondary antibodies were done for 90 min at room temperature in PBST. Washes in PBST were performed three times after primary and secondary antibody incubation, and incubations in phalloidin (1:200 dilution) in PBST were done for 15 min followed by wash at room temperature before mounting. Stained wings were mounted in 15 μl Vectashield mounting medium (Vector Laboratories). Wing discs were isolated from third instar larvae and submerged in fixation solution (4% paraformaldehyde in PBS at 4°) for 10 min. All other procedures were the same as for pupal wing immunohistochemistry.

Primary antibodies were as follows: mouse monoclonal anti-Fmi (1:200 dilution, DSHB), guinea pig polyclonal anti-Pk[C] (1:800 dilution, Olofsson et al., 2014), rat monoclonal anti-HA (clone 3F10, 1:200 dilution, Roche). Secondary antibodies from Thermo Fisher Scientific were as follows: 488-donkey anti-mouse, 488-goat anti-guinea pig, 546-donkey anti-goat, 633-goat anti-guinea pig, 633-goat anti-rat, 647-donkey anti-mouse. Alexa 635 and Alexa 350 conjugated phalloidin were from Thermo Fisher Scientific.

### Quantification of Vang::EYFP polarity

28hr APF pupal confocal wing images were stacked into a max-projected image and oriented as proximal to the left, anterior at the top. Non-overlapping polarity nematics were generated with the FIJI plugin ‘TissueMiner’ (Aigouy *et al*., 2010; Etournay *et al*., 2016) and overlayed on these images in a distinct color from the background image (in this case red). A custom-built MATLAB script allows the user to mark two regions of interests (ROIs). This script identifies the uniquely colored nematic lines within a given ROI by connected component analysis and outputs the *x^i^_min_*, *x^i^_max_*, *y^i^_min_*, *y^i^_max_*, pixel coordinates of each nematic *i* inside each ROI. The length *l*^*i*^ is then defined as

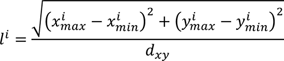

Where *d*_*xy*_ is the *xy* image resolution, and the angle ∠^*i*^ is defined as

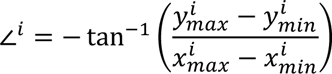

The MATLAB script (based on version R2021a with the Image Processing and Statistics and Machine Learning Toolboxes installed) has been deposited at https://github.com/SilasBoyeNissen/song2023.

### Western blot for Vang phosphorylation

Wing discs from wandering third instar larvae were rapidly dissected and transferred into ice cold PBS. Collected tissue was spun 3 minutes at 9,000g and the pellet snap frozen in liquid N2. 20-30 discs of each genotype were collected. To prepare extracts, 50 µL of lysis buffer with (all samples except *λ* phosphatase treatment) or without phosphatase inhibitors (sample for l phosphatase treatment) was added to thawed samples and the tissue lysed by multiple passages through a 27½ G needle and incubated on ice for 30 minutes. Samples were then spun at 20,000g for 15 minutes at 4°C, the supernatant collected, and protein concentration measure by Bradford.

### Lysis buffers

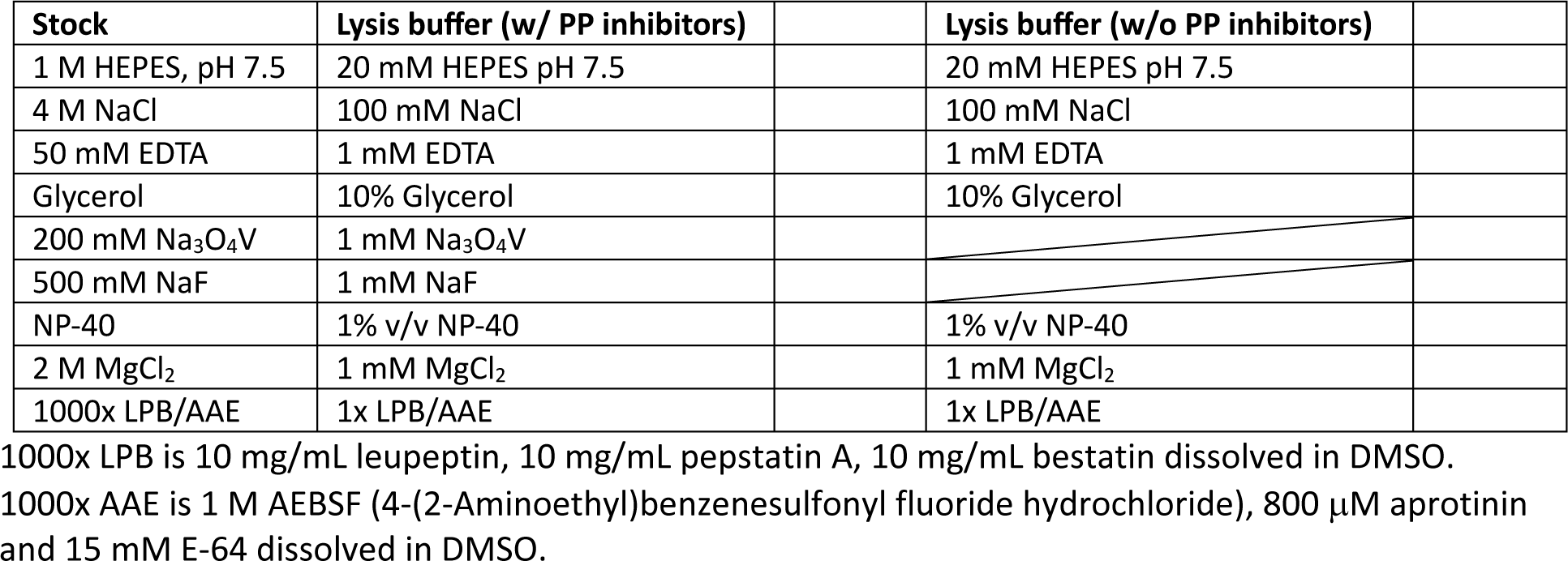

For *λ* phosphatase treatment, 2 µL of *λ* protein phosphatase (NEB P0753S) was added and incubated at 37°C for 30 minutes and then placed on ice. 20 µg total protein per lane was mixed with 4x LDS (Invitrogen) 200 mM DTT to a final of 1X LDS 50 mM DTT and incubated at 50°C for 15 minutes.

Samples were loaded and run in a 7% Tris-acetate gel (Invitrogen) in tris-acetate buffer on ice at 45V. Transfer to a pre-activated PVDF membrane was done in 10 % MeOH Towbin buffer at 4°C for 70 minutes at 400 mA constant intensity.

The membrane was cut at approximately 50KDa. The low molecular weight portion was blocked in 10% Seablock diluted v/v in TBST (Tris-buffered saline with 0.1% Tween^®^ 20 detergent) for 30 min, and the piece of membrane with larger molecular weight proteins in 5% milk diluted w/v in TBST.

Membranes were incubated overnight in primary antibody o/n at 4°C: Rabbit ⍺-Vang (Rawls & Wolff, 2003) diluted 1:1,000 in 5% milk in TBST or Rabbit ⍺-Actin (Sigma) diluted 1:2,000 in 10% Seablock in TBST followed by 3x wash in TBST for 10 minutes each. Membranes were developed in ECL reagent and imaged.

*Genotypes for flies in each Figure:*

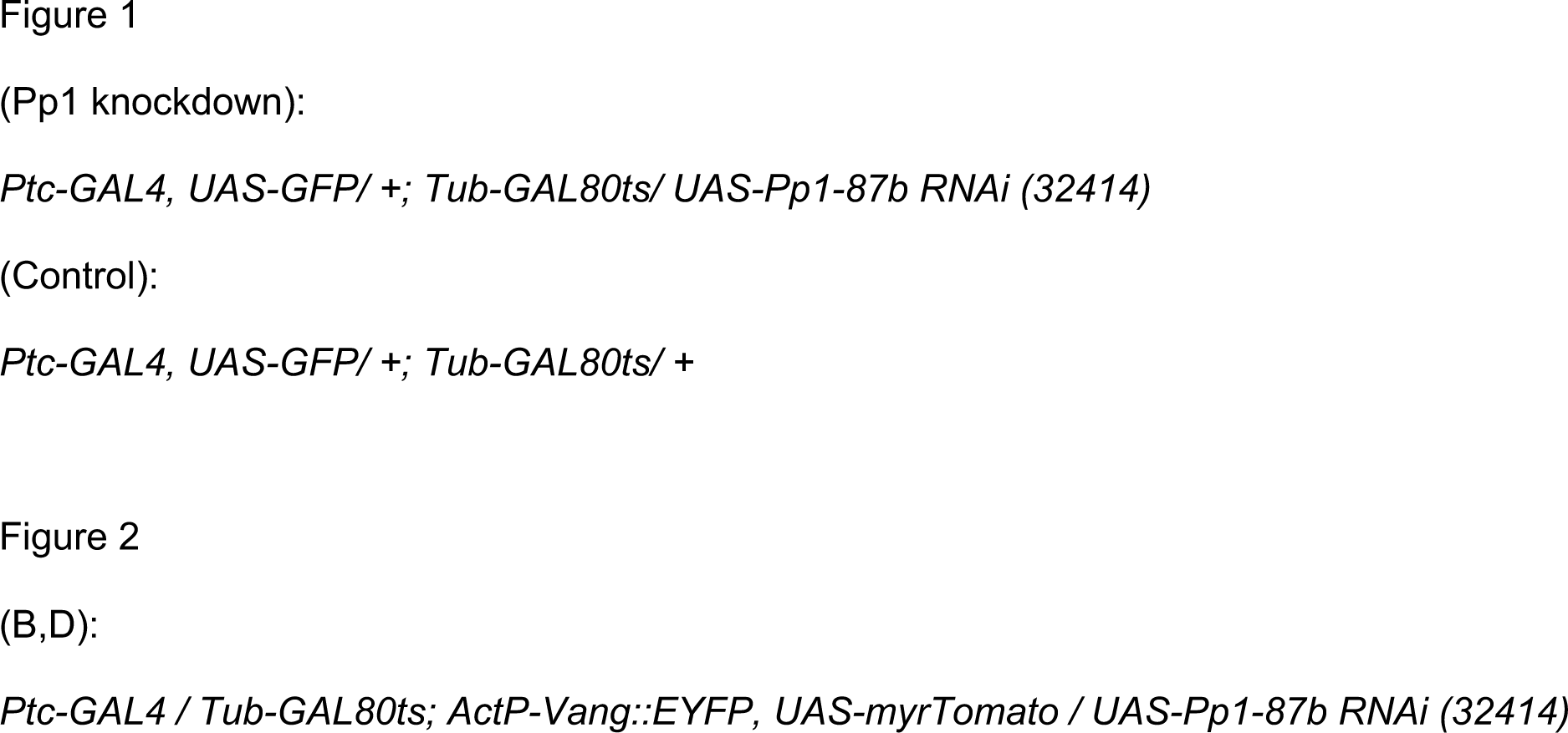
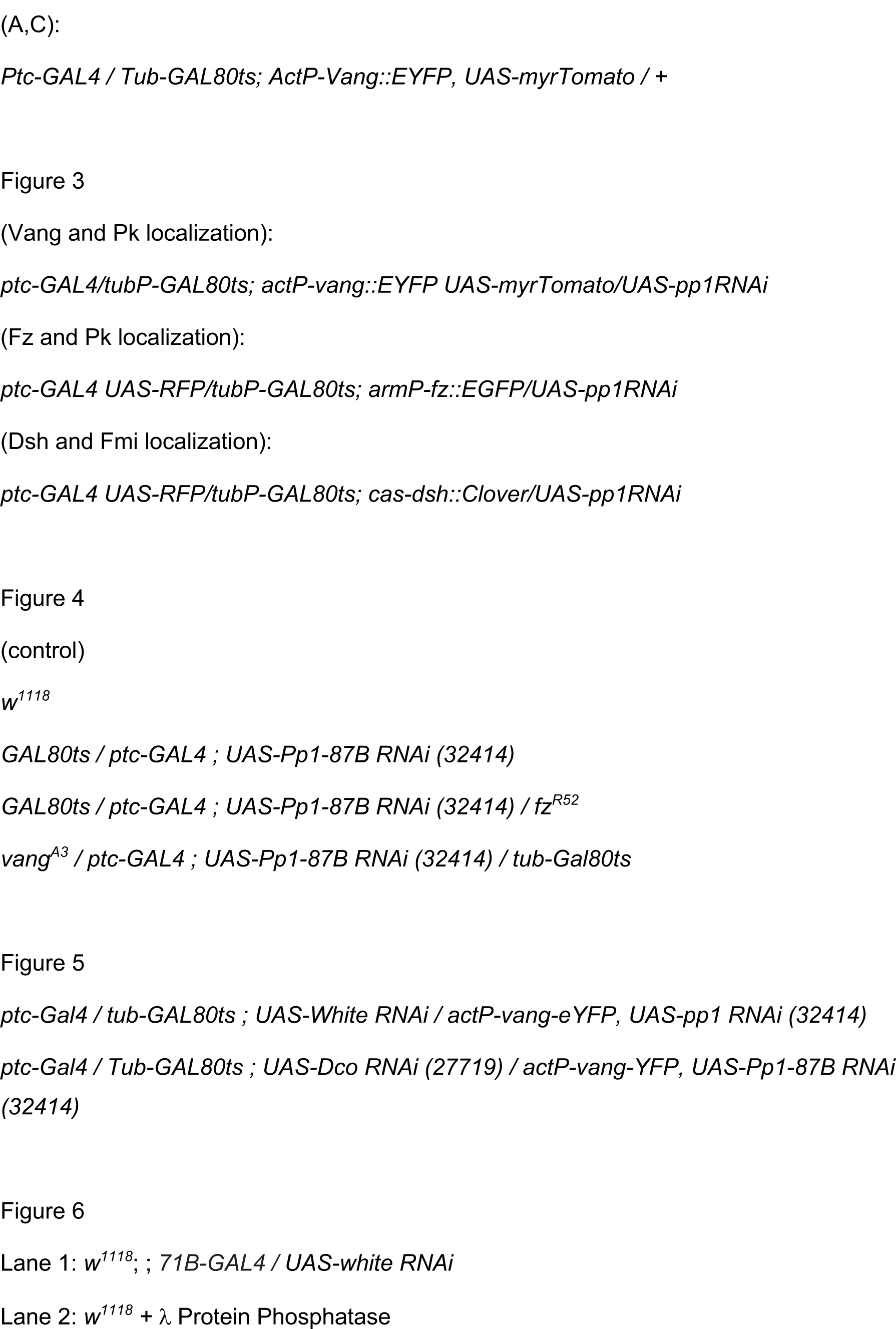
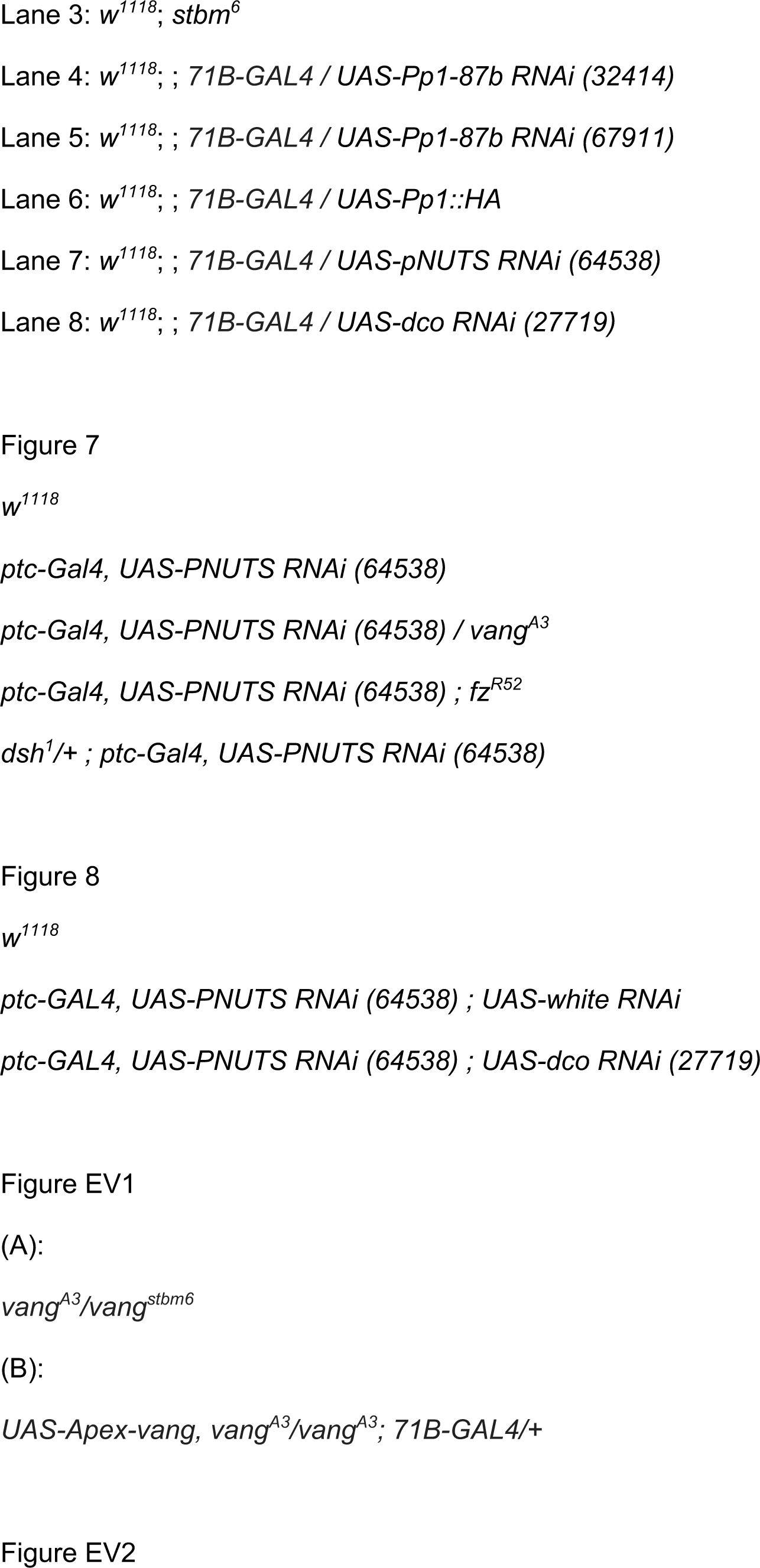
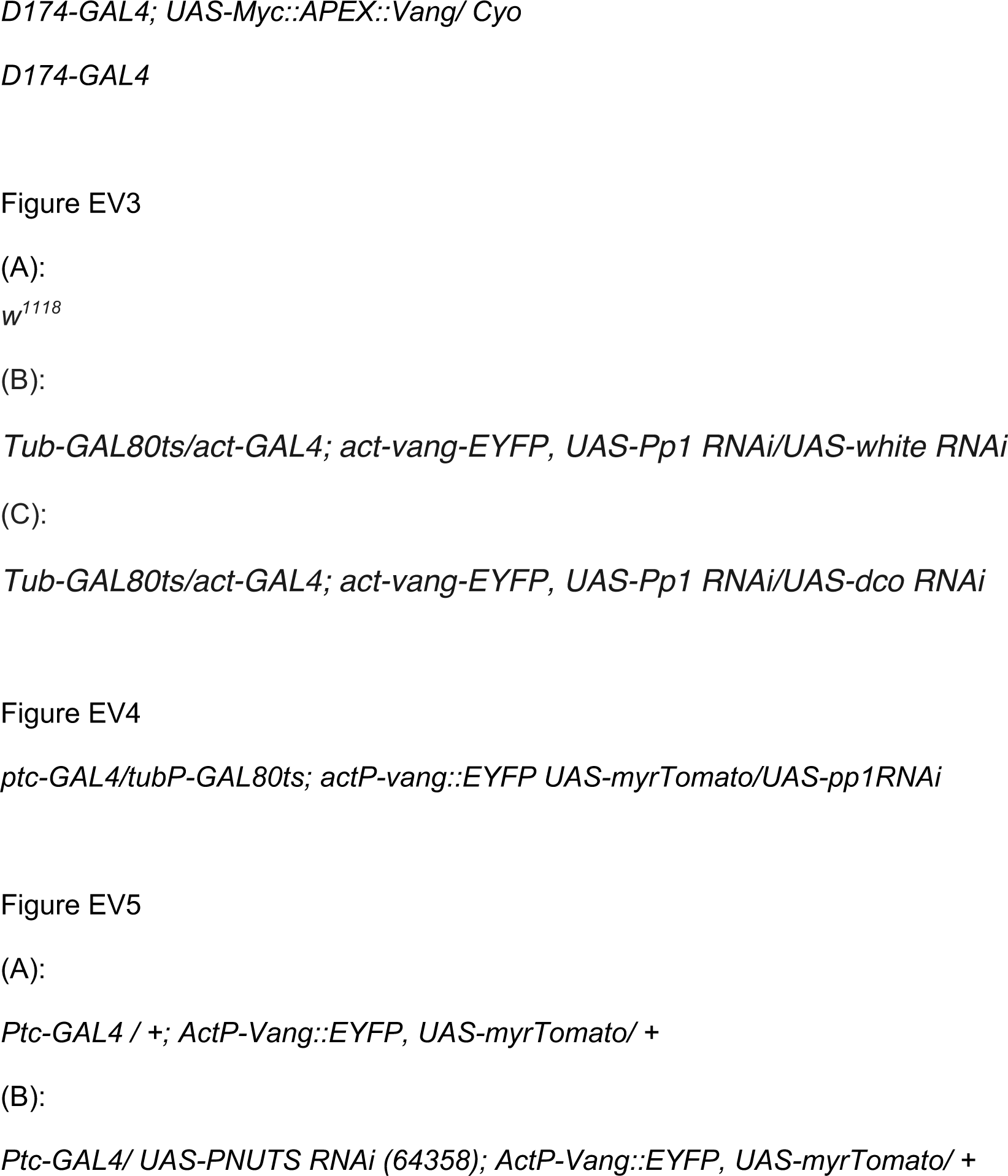

Drosophila stocks not maintained in public repositories are available upon request.

## Supporting information

Expanded View Datasets

## Acknowledgements

This work was funded by NIH R35GM131914 (J.D.A.). N.P. is an investigator of the Howard Hughes Medical Institute (HHMI). We acknowledge support from NIH Transformative R01 grant 5R01DK121409 (S.A.C. and N.P). S.B.N. was supported by the Novo Nordisk Foundation (grant awards NNF20OC0059462 and NNF21CC0073729) and the Stanford Bio-X Program.

## Data availability

RNAi screen results: RSVP database (https://www.flyrnai.org/cgi-bin/RSVP_search.pl)

Polarity measurement computer scripts: GitHub (https://github.com/SilasBoyeNissen/song2023)

---

**Expanded View Figure EV1.**
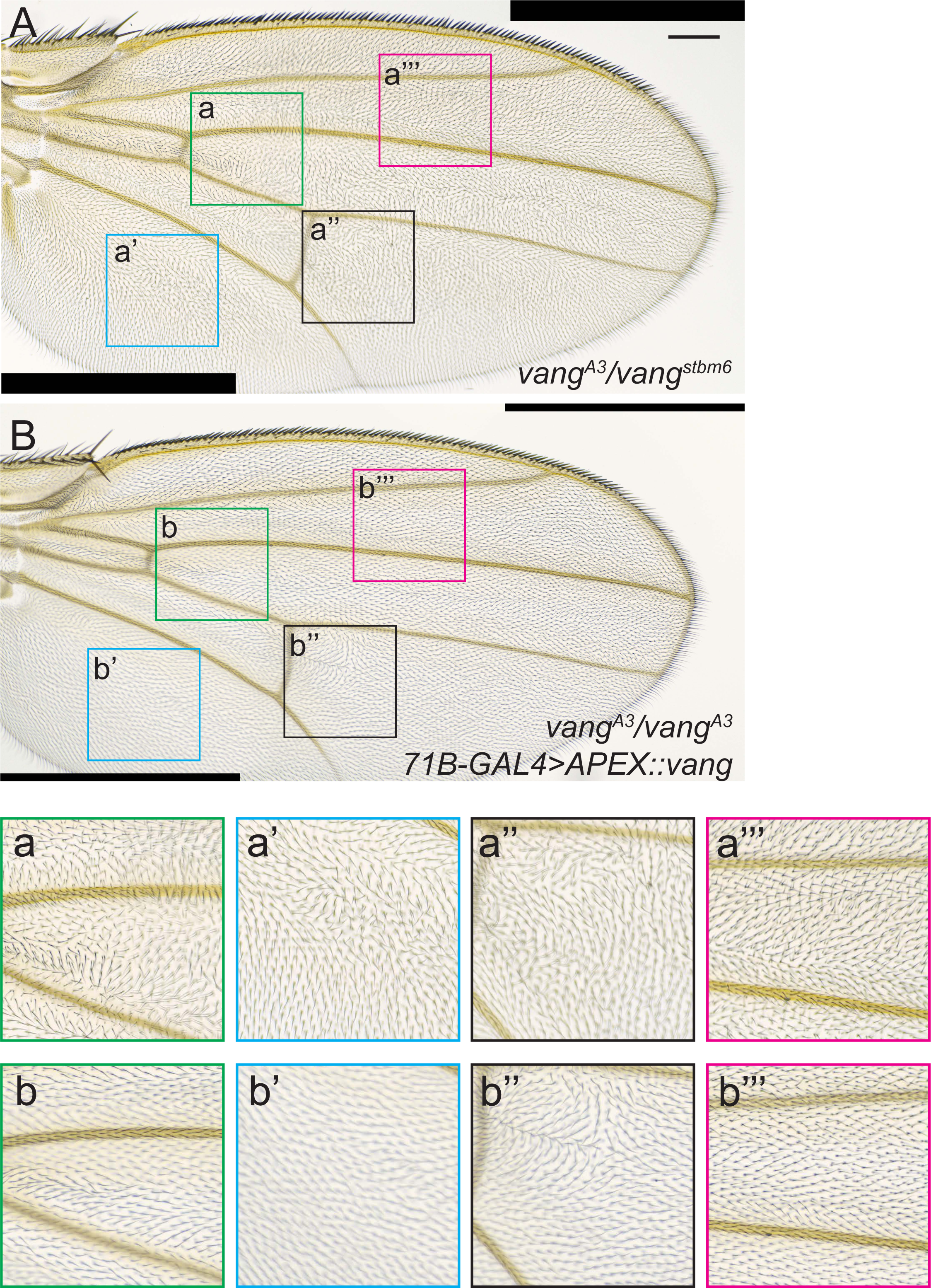
**Rescue of *vang^stbm6^***. Rescue of the *vang* mutant phenotype by APEX::Vang expression. A *vang* (*vang^A3^/vang^stbm6^*) mutant wing grown at 25° (**A**) shows the stereotypical PCP polarity phenotype, whereas a *vang* mutant wing ubiquitously expressing APEX::Vang (*UAS-Apex-vang, vang^A3^/vang^A3^; 71B GAL4/+*) grown at 25° (**B**) shows almost complete rescue. Magnified images of the boxed areas (a-a’’’ and b-b’’’) show nearly complete to complete rescue, with the exception of one area distal to the posterior crossvein (b’’) that typically showed disturbed polarity. This is likely an overexpression phenotype, as growth at 29° made this polarity disturbance worse. Scale bar = 100μm.

**Expanded View Figure EV2.**
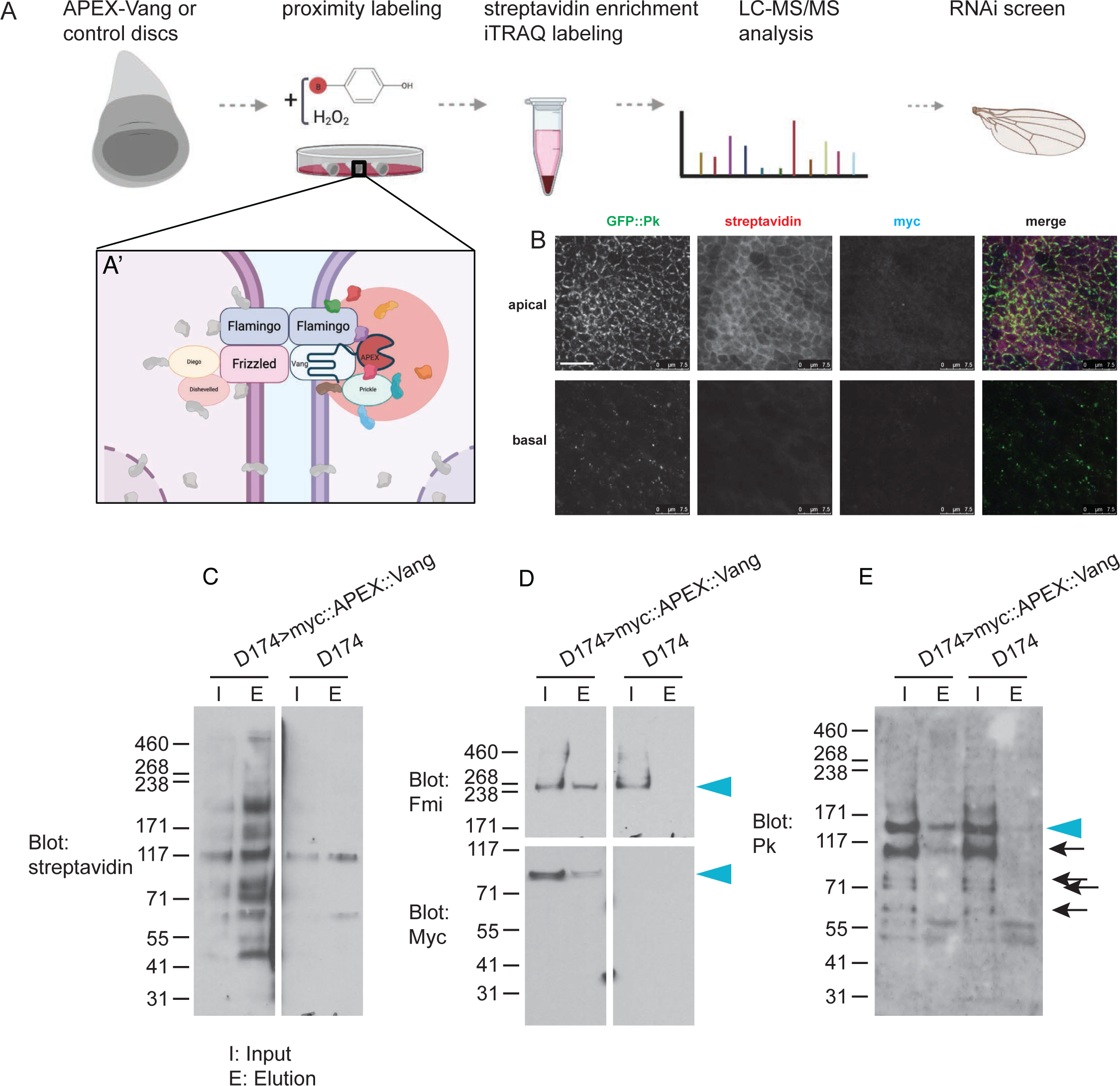
Schematic and validation of the APEX labeling protocol. **A-A’**. Workflow for proximity labeling. **A’**. Labeling is expected to biotinylate proteins in close proximity to the APEX::Vang probe. **B**. Labeled disc showing GFP::Pk, streptavidin detecting biotinylated proteins, Myc detecting Myc::APEX::Vang. **C**. Input and biotin enriched elution of APEX::Vang and control (D174) total protein. **D**. Input and elution probed for Fmi. The filter was cut and the upper portion probed for Fmi and the lower portion probed for Myc. Fmi and Myc::APEX::Vang are detected in the APEX::Vang but not the control elution (arrowheads). **E**. Input and elution probed for Pk. Pk is detected in the APEX::Vang but not the control elution (arrowhead). Degradation products of Pk are present in this blot (arrows). Scale bar = 7.5μm.

**Expanded View Figure EV3.**
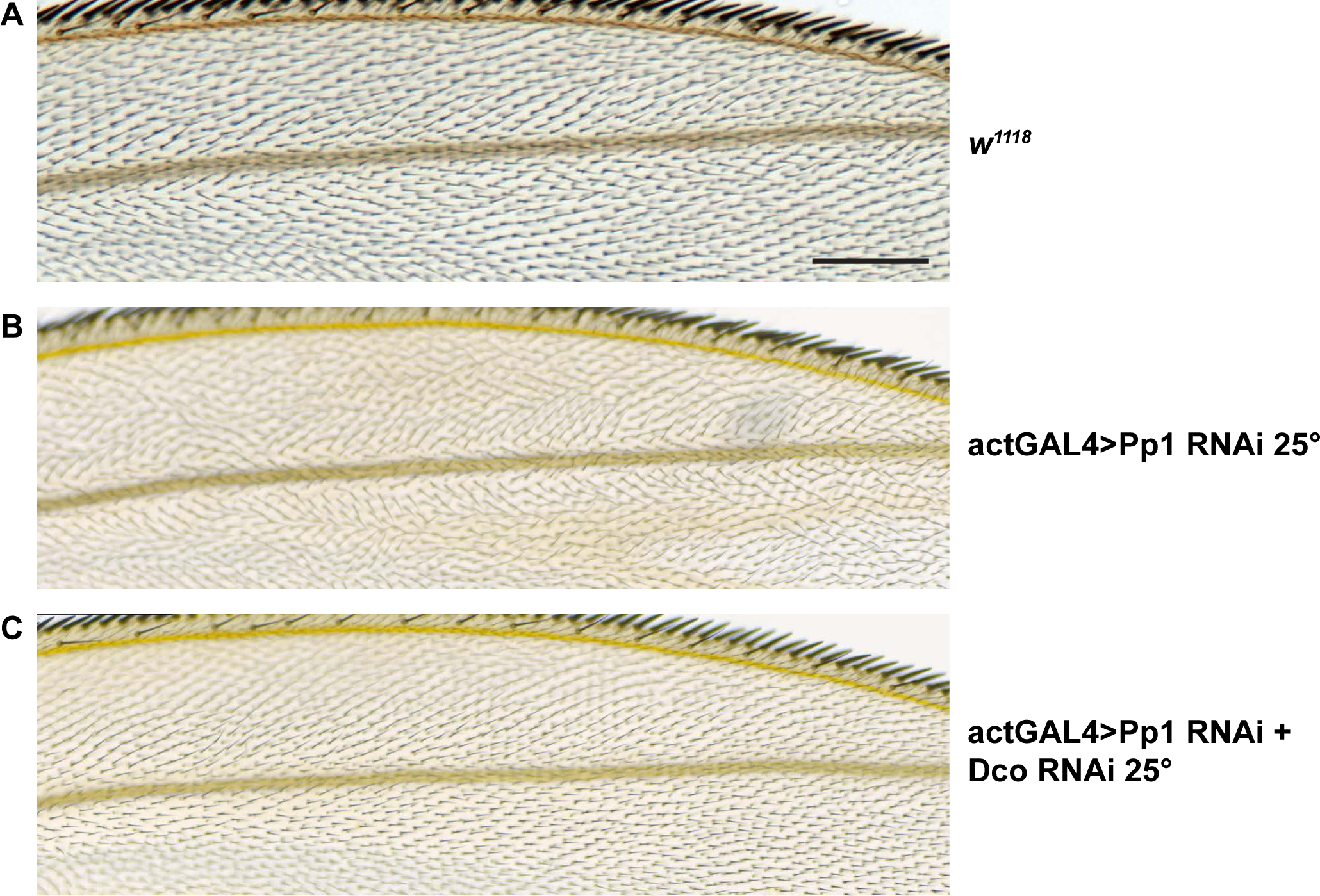
ActinGAL4 driving *Pp1-87B* RNAi throughout the wing induces polarity defects outside the *ptc* domain that are suppressed by simultaneous knockdown of Dco. **A.** Control (*w^1118^*) wing showing a region near the anterior wing margin. **B.** *Pp1-87B* (TRiP RNAi stock 32414) plus *wRNAi* knockdown raised at 25°C shows polarity disturbance in this region. **C.** Simultaneous expression of *Dco* RNAi and *Pp1-87B* (TRiP RNAi stock 32414) raised at 25°C suppresses the polarity defect. >10 wing for each of A-C were examined. Scale bar = 100μm.

**Expanded View Figure EV4.**
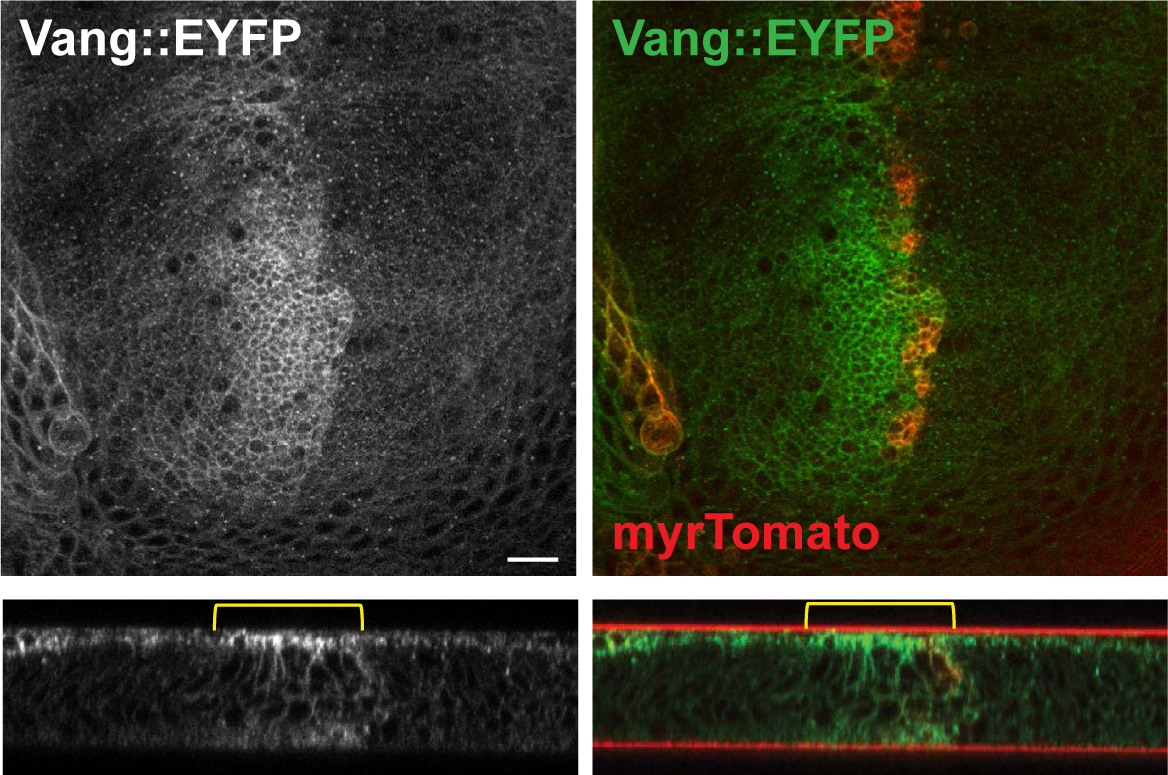
Vang accumulates at apical junctions and basolaterally in regions of a third instar wing disc where Pp1-87B is knocked down. XY and Y-Z planes are shown. myrTomato labels the *ptc* domain where *Pp1-87B* (TRiP 32414) RNAi is expressed. Scale bar = 10μm.

**Expanded View Figure EV5.**
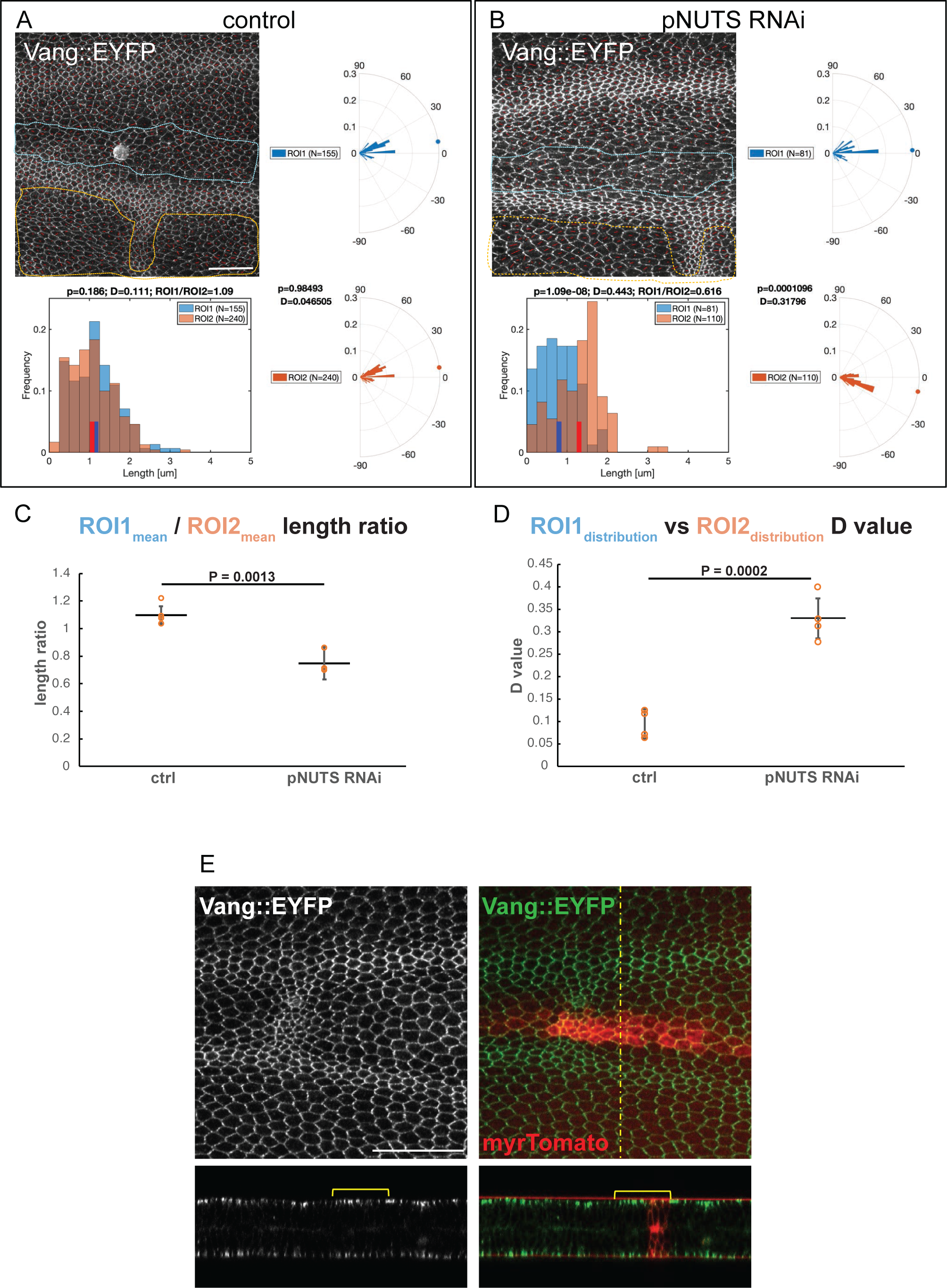
Knockdown of PNUTS in the *ptc* domain disrupts polarization of Vang subcellular localization. Knockdown (TRiP RNAi stock 64538) was driven by *ptc-GAL4* in wings expressing Vang::EYFP, and pupal wings were harvested at 28 hr APF. The degree of polarization and orientation of polarization of Vang::EYFP was compared for cells in the *ptc* domain, defined by mRFP expression (ROI1), and an adjacent domain posterior to the L4 vein where no knockdown occurred (ROI2). Polarization of individual cells was determined using the the Fiji plug-in “Tissue Analyzer” to determine nematics (red overlay lines), and both the length and orientation of each nematic was extracted using a custom MATLAB script (see Methods). Nematic lengths are plotted in histograms and orientations in rose plots. Individual examples are shown in **A,B**. N indicates the number of cells included in the region of interest (ROI). **C**. Ratios of mean nematic length, indicating degree of polarization in the *ptc* domain compared to the posterior domain, for four control and four PNUTS knockdown wings are plotted. PNUTS knockdown produced a significant decrease in mean length ratio (degree of polarization) (P = 0.0013, Students t-test). **D**. Distributions of polarity orientation for the *ptc* domain and the posterior domain were compared for the same four wings using the Kolmogorov-Smirnov test where D close to 0 indicates identical and D closer to 1 indicates different distributions. D values for PNUTS knockdown are significantly greater (P = 0.0002, Students t-test), confirming the visual impression that orientations are more dispersed in the *ptc* domain compared to the posterior domain. **E**. PNUTS knockdown does not cause excess accumulation or relocalization of Vang. Despite the diminution of polarization of Vang::EYFP upon PNUTS knockdown (Appendix Figure S4), Vang::EYFP does not accumulate to excess in apicolateral junctions or on the lateral membranes as was observed with knockdown of Pp1-87B (Figure 3). Scale bars = 50μm.

## Appendix

**Appendix Figure S1.**
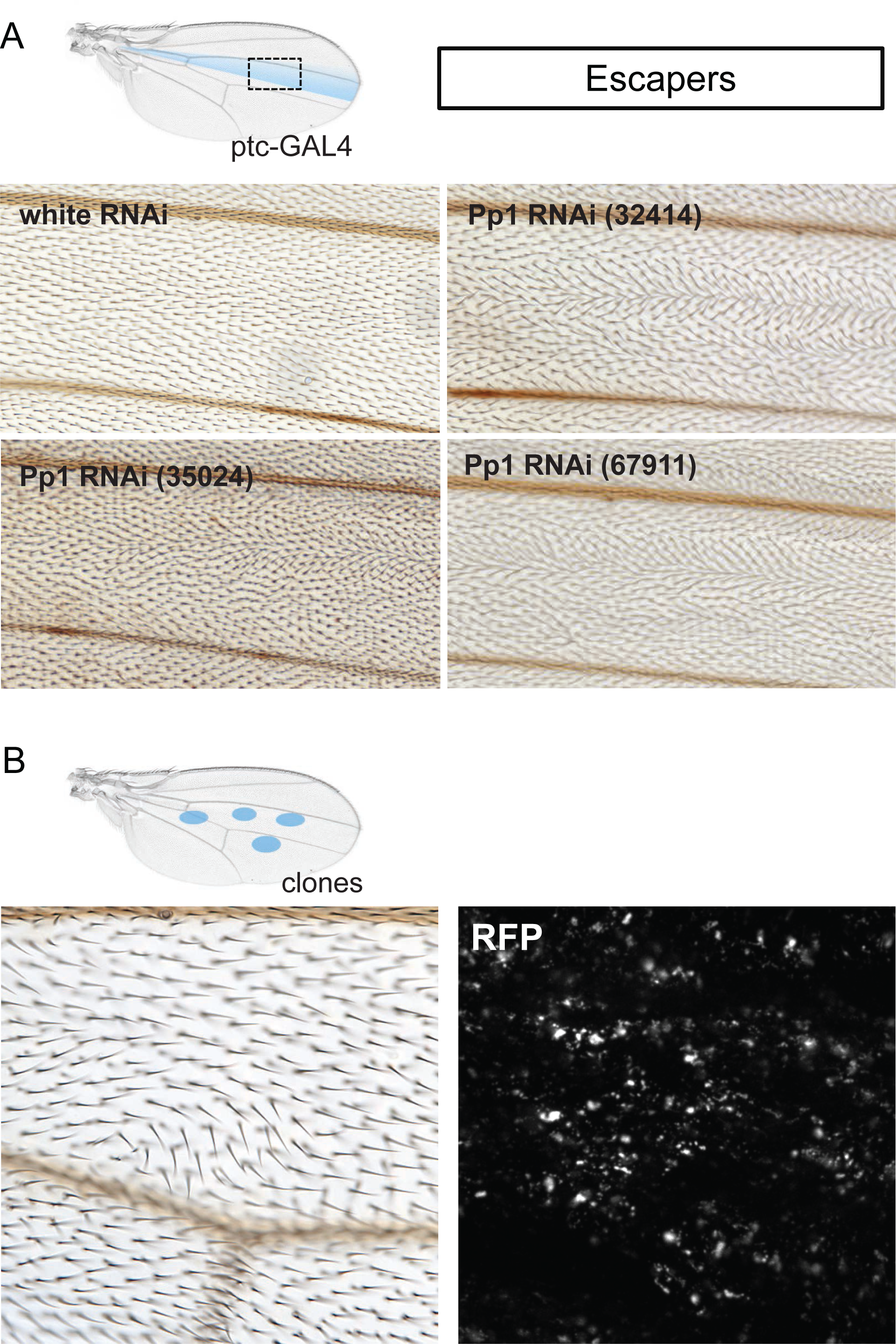
Wing hair polarity phenotype of escapers expressing three Pp1- 87B TRiP RNAi lines raised at 22°. **A**. Wings from escapers expressing control (*w^1118^*) or *Pp1-87B* RNAi in the *ptc* domain. **B**. Wing from a fly expressing *Pp1-87B* RNAi in clones. Clones were marked with RFP, but RFP in pupal wings was not detected in discrete clones, suggesting considerable amounts of cell death. Genotypes: (A): *UAS-Dcr-2/ +; ptc-GAL4/ +; UAS-white RNAi/ +* *UAS-Dcr-2/ +; ptc-GAL4/ +; UAS-Pp1-87b RNAi (32414)/ + UAS-Dcr-2/ +; ptc-GAL4/ +; UAS-Pp1-87b RNAi (35024)/ + UAS-Dcr-2/ +; ptc-GAL4/ +; UAS-Pp1-87b RNAi (67911)/ +* (B): *y w hsflp/ +; ActP>CD2>GAL4, UAS-RFP/ UAS-Pp1-87b RNAi (32414)/ +*

**Appendix Figure S2.**
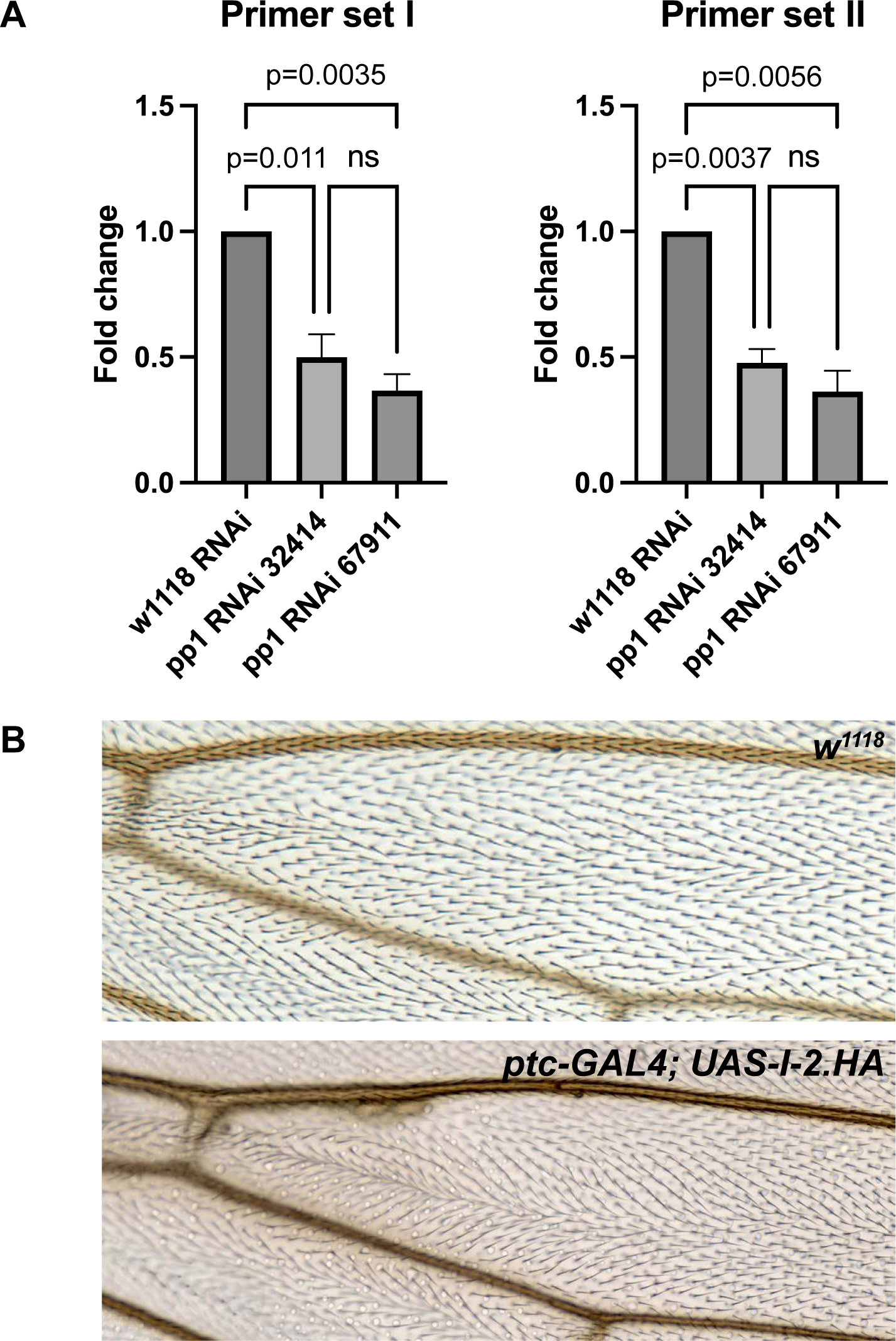
RNAi Validation. **A.** qRT-PCR was performed for two Pp1-87B RNAi lines, TRiP RNAi stock 32414 (BM32414) and TRiP RNAi stock 67911 (BM67911) driven by *71B-GAL4*, using two independent primer sets (I and II). In third instar discs, each reduced Pp1 RNAi levels by roughly half. Three biological replicates, each with three technical replicates, were performed (SEM is shown; p values as labeled, Student’s T-test). **B.** The Pp1-87B knockdown phenotype was phenocopied by expression of the Pp1-87B inhibitor, Inhibitor-2 (I-2.HA; BDSC24101) under *ptc-GAL4* control. The similar phenotypes of the three RNAi lines to each other and to that resulting from expression of the inhibitor provide strong evidence of specificity. Genotypes: (A): *UAS-white RNAi/ 71B-GAL4* *UAS-Pp1-87b RNAi (32414)/ 71B-GAL4 UAS-Pp1-87b RNAi (67911)/ 71B-GAL4* (B): *ptc-GAL4/ UAS-I-2.HA*

**Appendix Figure S3.**
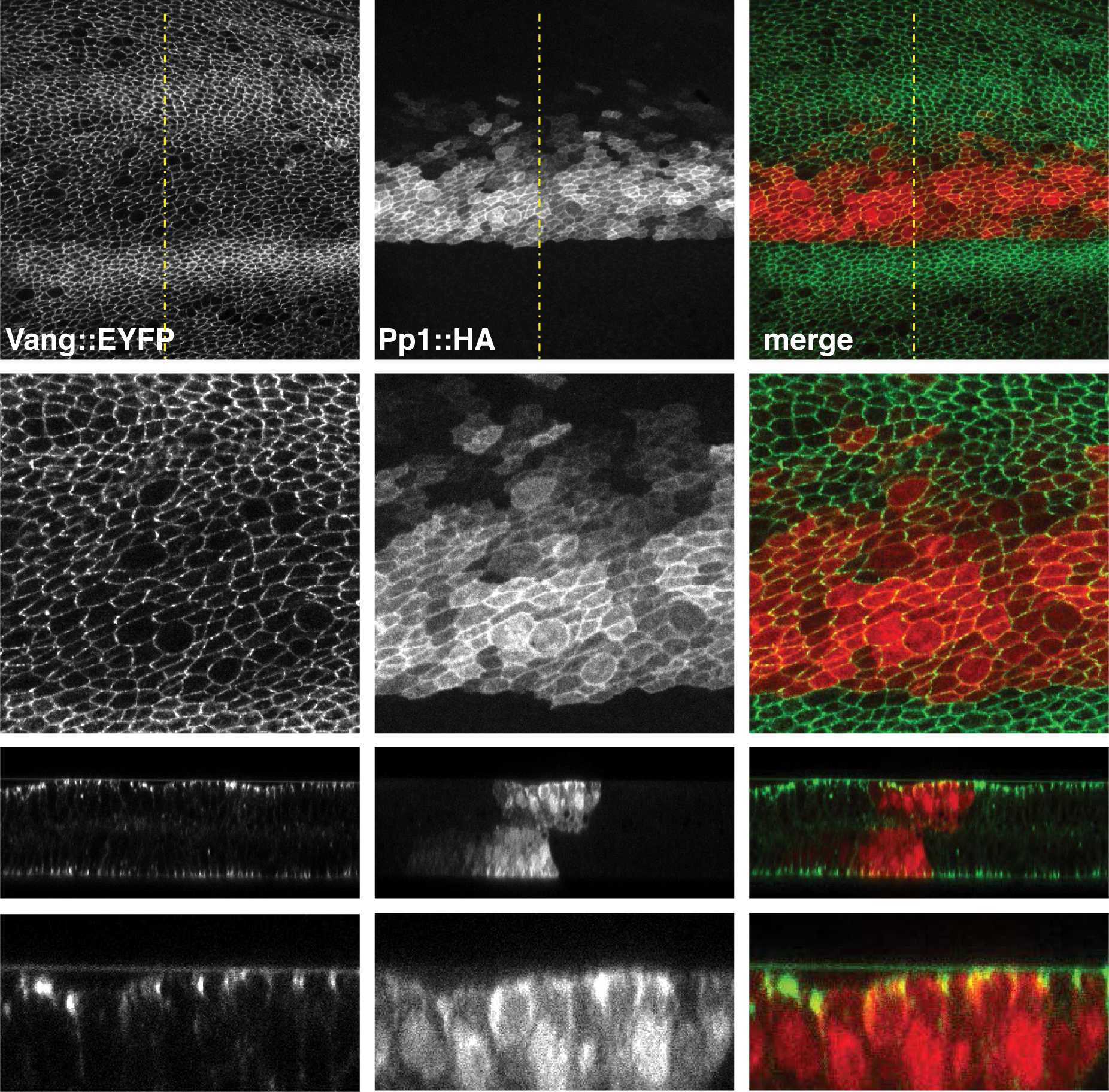
Pp1-87B is expressed apicolaterally and in the cytoplasm of pupal wings. 18 hr Vang::EYFP pupal wing in which Pp1-87B::HA is expressed in the *ptc* domain. EYFP and HA are shown in the XY and Y-Z planes. Apicolateral junctional expression of Pp1-87B is consistent with the possibility that it acts on core PCP proteins. Genotypes: *ptc-GAL4/actP-vang::EYFP; UAS-mCherry/UAS-pp1::HA*

**Appendix Figure S4.**
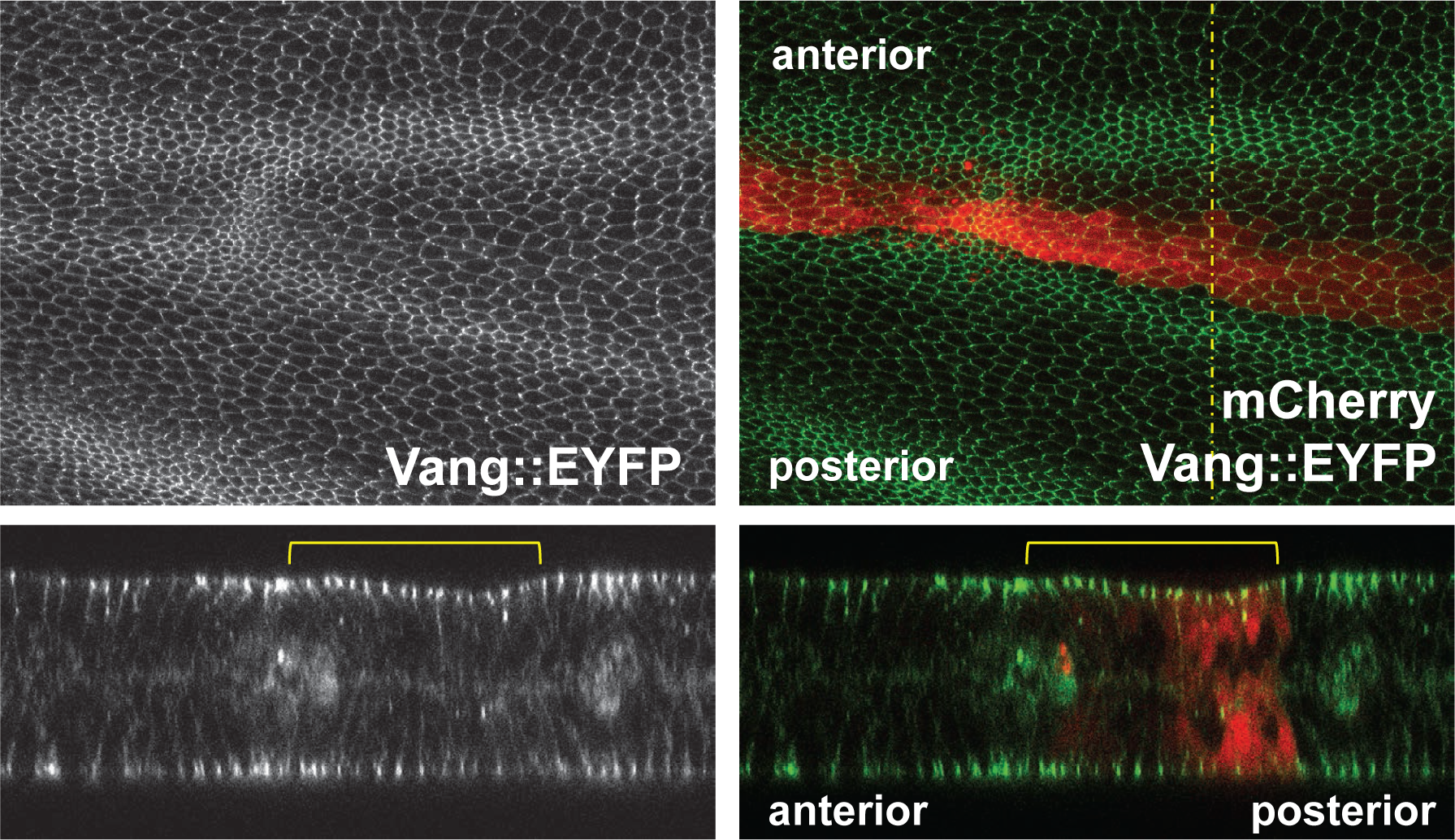
Expression of Pp1-87B in the *ptc* domain did not significantly modify Vang apical-basal localization. X-Y and Y-Z planes from a Vang::EYFP expressing wing ectopically expressing Pp1-87B::HA in the *ptc* domain show that Pp1- 87B::HA expression does not appreciably alter Vang::EYFP localization. Genotypes: *ptc-GAL4/actP-vang::EYFP; UAS-mCherry/UAS-pp1::HA*

**Appendix Figure S5.**
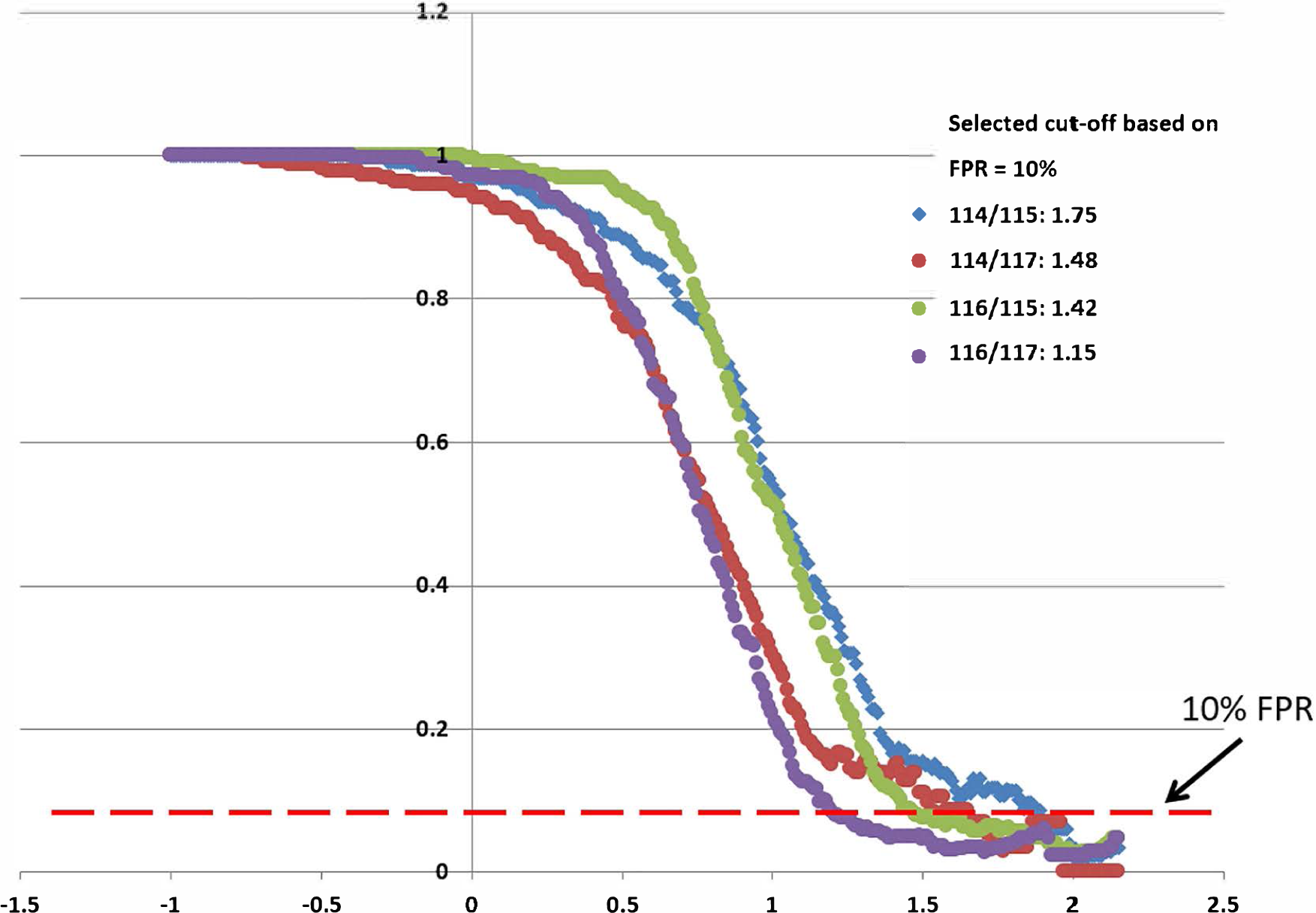
Determination of the Cutoff Point for Proteome Analysis. 114 and 116 are the experimental samples while 115 and 117 are the control samples. The iTRAQ ratios of experimental sample over control samples were calculated as 114/115, 114/117,116/115 and 116/117 respectively. Genes on the positive control (PC) list were selected manually from literature while the transcription factors and metabolic enzymes are assembled as negative control (NC). PC genes usually have higher iTRAQ ratios compared to NC genes. The threshold of the iTRAQ ratio is determined based on a false positive rate of 0.1 (dashed line), which means that a protein is 10 times more likely to be a true positive protein than a false positive.

